# STAT1 is required to establish but not maintain IFNγ-induced transcriptional memory

**DOI:** 10.1101/2022.09.17.508285

**Authors:** Sahar S.H. Tehrani, Pawel Mikulski, Izma Abdul-Zani, João F. Mata, Wojciech Siwek, Lars E.T. Jansen

## Abstract

Exposure of human cells to interferon-γ (IFNγ) results in a mitotically heritable yet reversible state called long-term transcriptional memory. We previously identified the clustered GBP genes as strongly primed by IFNγ. Here we discovered that in primed cells, both interferon-responsive transcription factors STAT1 and IRF1 target chromatin with accelerated kinetics upon re-exposure to IFNγ, specifically at promotors of primed genes. Priming does not alter the degree of IFNγ-induced STAT1 activation or nuclear import, indicating that memory does not alter upstream JAK-STAT signalling. We found STAT1 to be critical to establish transcriptional memory but in a manner that is independent of mere transcription activation. Interestingly, while Serine 727 phosphorylation of STAT1 was maintained during the primed state, STAT1 is not required for the heritability of GBP gene memory. Our results suggest that memory of interferon exposure constitutes a STAT1-mediated, heritable state that is established during priming. This renders GBP genes poised for subsequent STAT1 and IRF1 binding and accelerated gene activation upon a secondary interferon exposure.

## Introduction

The innate immune system, in contrast to adaptive immunity, has classically been considered non-specific and transient with no memory of prior infections. However, it has become apparent that in some cases, activation of an innate immune response can lead to a primed state that results in enhanced resistance to reinfection, even to a different pathogen (Netea, Quintin, and Van Der Meer 2011; Peignier and Parker 2020). Such priming can last for weeks or months, is reported to be independent of the adaptive immune system and is referred to as trained immunity (Netea, Quintin, and Van Der Meer 2011). Examples include exposure of mice that lack functional B and T lymphocytes to BCG vaccine (Bacillus Calmette–Guérin), *Candida albicans*, or β-glucan (a component of the fungal cell wall) that induce a primed response resulting in enhancement of inflammatory cytokine production upon a secondary infection (Kleinnijenhuis et al. 2012, Quintin et al. 2012). This innate immune priming correlates with molecular changes including chromatin accessibility and modification (Lau et al. 2018), transcription of long non-coding RNAs (lncRNAs) (Fanucchi et al. 2019), DNA methylation (Verma et al. 2017) and reprogramming of cellular metabolism (Natoli and Ostuni 2019; Netea et al. 2020). At present, most of these molecular signatures are correlative, yet understanding the mechanistic basis of “trained immunity” would enable us to exploit this phenomenon for clinical applications such as vaccination as well as for the prevention and treatment of conditions such as chronic inflammation.

While the molecular drivers of trained immunity remain elusive and correlative, an analogous priming effect can occur at the level of gene expression that may contribute to trained immunity. This effect, called long-term transcriptional memory is observed upon exposure to inflammatory cytokines such as TNF-α and interferons even in non-immune cells (Gialitakis et al. 2010; Zhao et al. 2020; Light et al. 2013). Transcriptional memory is also observed outside of the mammalian immune system in a variety of species ranging from yeast to plants, allowing organisms to adapt faster to previously encountered environmental stress conditions such as nutrient deprivation (‘D’Urso and Brickner 2017), heat (Ding et al. 2013; Lämke et al. 2016) and cold stress (Song et al. 2012).

Possible mechanisms of transcriptional memory can be broadly categorized as “*cis*-acting” constituting factors such as DNA or histones modification which may be locally inherited through the cell cycle (Moazed 2011; Quintin et al. 2012); and ”*trans*-acting” such as soluble transcription factors that initiate and/or maintain memory of the signal even in its absence, e.g. through rewiring of signalling cascades or transcription factor networks (Moazed 2011).

Several lines of evidence indicate that local cis-regulated factors can contribute to memory. For instance, DNA demethylation has a positive impact on TNF-α-mediated transcriptional memory genes (Zhao et al. 2020). Additionally, dimethylation of histone H3 on lysine 4 (H3K4me2) has been widely associated with the maintenance of a prime state. For instance in yeast, COMPASS and mediator play a role in maintaining of H3K4me2 in the context of memory of INO1 expression (‘D’Urso et al. 2016). In plants, both H3K4me2 and 3 are implicated in retaining transcriptional memory of a prior stressor such as acquired thermotolerance (Lämke et al. 2016). Beside a role for local chromatin factors in transcriptional memory, transacting transcription factors have also been implicated in initiation or maintenance of priming. The yeast transcription factors, Sfl1 and Tup1 are critical for maintaining poised transcription and loss of those factors disrupts transcriptional memory of INO1 and GAL1, respectively. (D’Urso et al. 2016; Sood et al. 2017). Moreover, transcription factor MYC2, which is induced upon dehydration stress and HSFA2 for heat stress are required for memory (Liu and Avramova 2016; Lämke et al. 2016).

The widespread occurrence of transcriptional memory suggests that some basic principles and mechanisms may underlay this phenomenon and may thus be driven by mechanisms that are not unique to one cell type or system. In the context of the innate immune system, it is striking that cytokine signals such as interferons induce transcriptional memory even in non-immune cells (Gialitakis et al. 2010). This creates an opportunity to discover general principals of transcriptional memory underlying trained immunity without the confounding effects of immune signalling and cell differentiation that can be induced by cytokines.

A well-established paradigm in innate immune transcriptional memory is the priming of genes by interferon-γ (IFNγ). In this case, a subset of IFNγ-activated genes can maintain a heritable poised state in the absence of active transcription of the primed genes. Yet, the primed state leads to an enhanced expression upon re-exposure to IFNγ (Figure 1A). Early studies showed that an IFNγ target gene, HLA-DRA undergoes priming which correlates with the maintenance of RNA polymerase II (RNA PolII) (‘D’Urso and Brickner 2017; Light et al. 2013) and H3K4me2 on the promoter of HLA-DRA in primed cells (Gialitakis et al. 2010), at least short term up to 48 hours post an IFNγ pulse. However, other reports did not detect RNA Pol II poising following priming of mouse fibroblasts by IFNβ and IFNγ or HeLa cells (Kamada et al. 2018; Siwek et al. 2020). Additionally, H3.3 and H3K36me3 were observed as a memory mark, maintained on primed genes, albeit for a short 2-day period following an IFNγ pulse (Kamada et al. 2018; Siwek et al. 2020).

**Figure 1.**
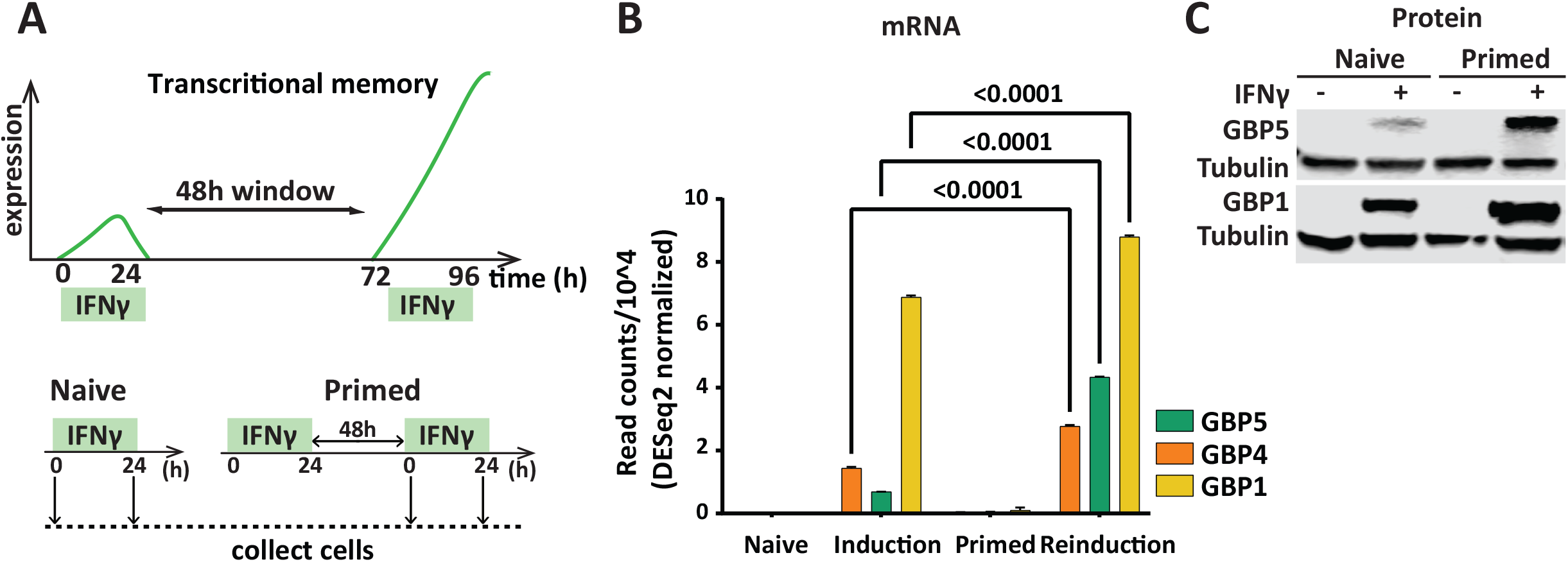
Long-term transcriptional memory of GBP genes (A) Top: Principle of IFNγ induced transcriptional memory. Among IFNγ inducible genes, those with memory show faster and stronger expression upon a second induction with IFNγ. Bottom: Experimental outline for transcriptional memory; HeLa cells were primed with IFNγ for 24 hours, followed by IFNγ washout. After 48h, naïve and primed cells were induced by IFNγ for 24 h. **(B)** RNA-seq data showing transcriptional memory for GBP1, 4 and 5 genes [obtained from dataset reported in (Siwek et al 2020)] **(C)** HeLa cells were harvested at indicated time points and processed for western blotting probed for GBP1 and 5 protein levels, α-Tubulin (Tubulin) as loading control.

We previously reported that genes showing strong interferon-γ induced transcriptional memory tend to reside in genomic clusters and that the establishment of long-term memory of these genes is locally restricted by cohesion (Siwek et al. 2020). Here we aim to understand the role of transcription in priming and explore the contribution of the STAT and IRF transcription factors in IFNγ-mediated transcriptional memory. We find that in the primed state, the kinetics of the upstream JAK-STAT signalling cascade to activate STAT1 is not altered. Instead, we find that the chromatin of memory target gene promoters is more accessible in primed cells and that STAT1 and IRF1 are recruited faster specifically at the primed GBP cluster. Interestingly, memory is not driven by target gene transcription but relies upon a STAT1-dependent action during priming, after which STAT becomes dispensable for the maintenance of the primed state.

## Results

### GBP genes show long-term transcriptional memory

Exposure of cells to interferon-gamma (IFNγ) leads to a heritable, primed state resulting in enhanced activation of target genes following a second exposure (Gialitakis et al. 2010). Using this principle as an assay (outlined in Figure 1A), we previously identified genes encoding the guanylate binding proteins, including GBP1, GBP4 and GBP5 that show mitotically stable memory that is propagated for at least a week in proliferating cells [(Figure 1B) and (Siwek et al. 2020)]. Using a standardized protocol with a 2-day memory window (Figure 1A), we validated these findings by directly measuring protein levels for GBP1 and GBP5 in HeLa cell lines (Figure 1C). In this study, we use GBP genes as a readout of memory, particularly GBP5, 4 and 1 as they showed the strongest reinduction upon a second IFNγ exposure.

### Transcription of GBP1 is not sufficient to induce a local prime state

We previously reported that, based on single-cell RNA sequencing that, at least for GBP5, priming is manifested by an increased probability of primed cells to engage in target gene expression, correlating with the strength of the initial GBP5 activation (Siwek et al. 2020). Furthermore, earlier work has shown that priming results in enhanced Pol II recruitment or retention of promoter-bound polymerase in the absence of ongoing transcription (Kamada et al. 2018; Light et al. 2013). This suggests that transcription of the target gene itself may be sufficient to induce the prime state regardless of upstream signalling. To test this hypothesis directly, we artificially forced GBP1 transcription using CRISPR/Cas9 Synergistic Activation Mediator (CRISPRa-SAM) (Figure 2A), a method previously used for activation of a variety of genes in different cell types (Chavez et al. 2016; Konermann et al. 2014). In this way, we bypass the need for IFNγ and can determine the role of transcription in gene priming. We successfully established conditions for the CRISPRa-SAM activation of GBP1. A combination of 6 gRNAs targeting the GBP1 promotor, but not gRNAs for the unrelated ASCL1 control gene, is sufficient for GBP1 activation, as validated by RT-qPCR (Figure 2B).

**Figure 2.**
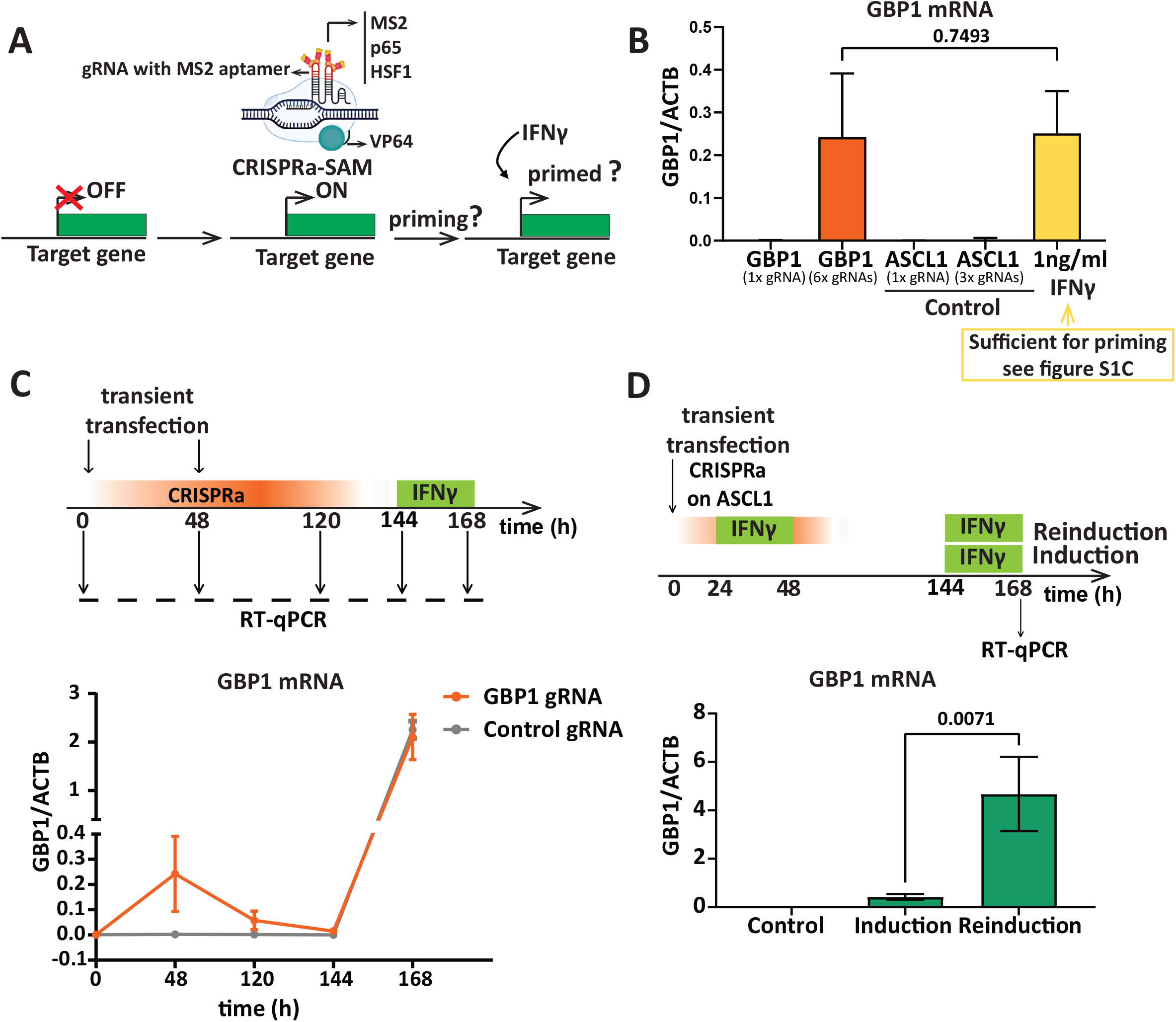
Transcription of a memory gene is not sufficient to induce a primed state. (A) Experimental outline; targeting of CRISPR/Cas9 Synergistic Activation Mediator (CRISPRa-SAM), results in forced gene activation, followed by IFNγ to determine priming state. **(B)** HeLa cells were transfected with dCas9-SAM technology with gRNA for GBP1 (memory gene) and ASCL1 (control). In parallel, cells were induced with 1 ng/mL of IFNγ for 24h for comparison. GBP1 mRNA expression was measured by RT-qPCR after 48h of transfection and normalized to ACTB expression. Error bars, SD; n=3. **(C)** Top: Experimental outline of CRISPRa targeting of GBP1 and ASCL1 (control) by two rounds of transient transfection with plasmids containing dCAS9-SAM, followed by induction with IFNγ. Bottom: RNA was isolated, GBP1 mRNA level was determined as indicted in C, by RT-qPCR and normalized to ACTB mRNA level. Data are shown as mean. Error bars, SD; n=3. **(D)** Top: Experimental outline; HeLa cells were transfected with dCas9-SAM with ASCL1 gRNA (control), after 24h cells were primed with IFNγ, followed by IFNγ washout. After 48h, naïve and primed cells were induced by IFNγ for 24h then cells were harvested after induction and reinduction. Bottom: GBP1 mRNA expression was measured by RT-qPCR after induction and reinduction, normalized to ACTB mRNA. Data are shown as mean. Error bars, SD; n= 3.

Titration of the concentration and duration of IFNγ exposure revealed that a 24h treatment of cells with only 1ng/mL of IFNγ is sufficient to activate GBP1 (Figure 2B, S1A, B) and induce a primed state that is heritable for at least 48h (Figure S1A, C). This level of IFNγ-mediated GBP1 mRNA is comparable to the level of induction by CRISPRa-SAM (Figure 2B). To test whether GBP1 expression *per se* is sufficient for priming we transfected HeLa cells with CRISPRa-SAM, targeted by either gRNAs specific for GBP1 or a control gene (ASCL1). We allowed CRISPRa-SAM-driven GBP1 expression to build up for 48h and then allowed cells to dilute out, prior to induction with IFNγ as outlined in Figure 2C. Our results demonstrate that GBP1 is not primed by prior CRISPRa-SAM activation despite similar transcriptional output as for IFNγ induction (Figure 2B). This indicates that mere transcription of GBP1 is not sufficient to establish transcriptional memory. Additionally, we confirmed that the CRISPRa-SAM procedure itself does not interfere with IFNγ-induced transcriptional memory by priming cells with IFNγ during CRISPR activation with a control gRNA (ASCL1) followed by reinduction with IFNγ (Figure 2D). Overall, unexpectedly, our results indicate that the recruitment of Pol II and the activation of the general transcription machinery is not sufficient to induce a prime state and that IFNγ signalling and its downstream transcription factors are necessary to initiate priming.

### Increased promoter accessibility of GBP4 and GBP5 in primed cells

As mere transcription does not appear to be the initiator of priming, we reasoned that IFNγ-specific transcription factors upstream of transcription initiation may be required for inducing long-term memory. We started out by determining promoter accessibility as an indirect readout of the degree of transcription factor binding and target gene activation during induction, memory and reinduction. We performed ATAC-seq in naïve and primed HeLa cells after inducing with IFNγ for 0h, 1h and 3h (Figure S2A). We find that the promoters of GBP5, and the adjacent GBP4 gene are selectively opened during IFNγ activation (Figure S2B). Interestingly, the GBP1 promoter is already in an accessible state in naïve cells and is not significantly opened further by IFNγ activation (Figure S2B). Moreover, in primed cells the GBP5 promoter and to lesser extent GBP4, shows accelerated opening during reinduction, particularly after 1 hour of IFNγ, but not at IFNγ target genes that do not show priming such as IRF1 and TAP1 [(Figure S2B) and (Siwek et al. 2020)].

In agreement with our previous report (Siwek et al. 2020) there is no indication of maintenance of an IFNγ-opened nearby enhancer site in primed cells, indicating that latent enhancers are not driving the accelerated reactivation of the GBP genes (Figure S2C).

### STAT1 and IRF1 are required for GBP5 expression

The increased ATAC signals at GBP promoters upon IFNγ induction suggest specific transcription factors target these promotors that may play a role in transcriptional memory. To explore this further, we first examined the role of transcription factors effectors of this pathway. The STAT and IRF family of proteins are the main transcription factors responding to interferons (Mogensen 2018). There are seven members in the STAT family (STAT1, 2, 3, 4, 5A, 5B and 6) and nine in the IRF family (IRF1-9) which target genes in response to different cytokines (Delgoffe and Vignali 2013; Antonczyk et al. 2019). STAT1 is well established as the key transcription factor in IFNγ signalling (Antonczyk et al. 2019). Upon stimulation, STAT1 is activated by the IFNγ receptor-bound JAK kinase. Phosphorylation results in homodimerization or heterodimerization with other STATs, leading to translocation into the nucleus and target gene activation (Rawlings, Rosler, and Harrison 2004). Genes encoding IRF transcription factors are activated by STAT1 that then cooperate with it to further induce downstream interferon-target genes (Schroder et al. 2004). One possible way of achieving a primed state is for a specific transcription factor to respond to IFNγ stimulation in a feedforward fashion. In such a scenario, IFNγ stimulation results not only in transcription factor activation but also in its continued expression, even after the removal of the cytokine. To determine whether any of the STATs or IRFs behave like this, we mined our RNA-seq dataset (Siwek et al. 2020) to determine their expression after induction, when primed and in the reinduction state (Figure 3A). As expected, several STAT and IRF members were strongly induced by IFNγ but following washout, all returned to baseline levels (Figure 3A). Thus, a model in which the key drivers of IFNγ-mediated gene expression engage in self-propagating expression, is unlikely. However, while the mRNA levels of these genes return to baseline, they may nevertheless be required to establish and/or maintain the primed state upon IFNγ exposure. To dissect the putative role of STAT and IRF proteins in priming in more detail in the HeLa model, we generated CRISPR/Cas9 knockout cell lines of a representative set of transcription factors. These include those most strongly activated by IFNγ; STAT1, STAT2, STAT3 and IRF1 as well as STAT5B and IRF9 (Figure S3A). Consistent with earlier reports, GBP5 induction is lost in STAT1 and IRF1 knockout cells [(Figure 3B) and (Ramsauer et al. 2007)], while STAT2, STAT5B and IRF9 are dispensable both for induction (Figure S3B) as well as priming (Figure S3C). Interestingly, STAT3 depletion had the opposite effect, leading to an increase in GBP5 induction (Figure S3B). From this analysis, we conclude that both STAT1 and IRF1 are required for GBP5 induction. Next, we asked if these two key transcription factors have any role in establishing or maintaining GBP5 priming.

**Figure 3.**
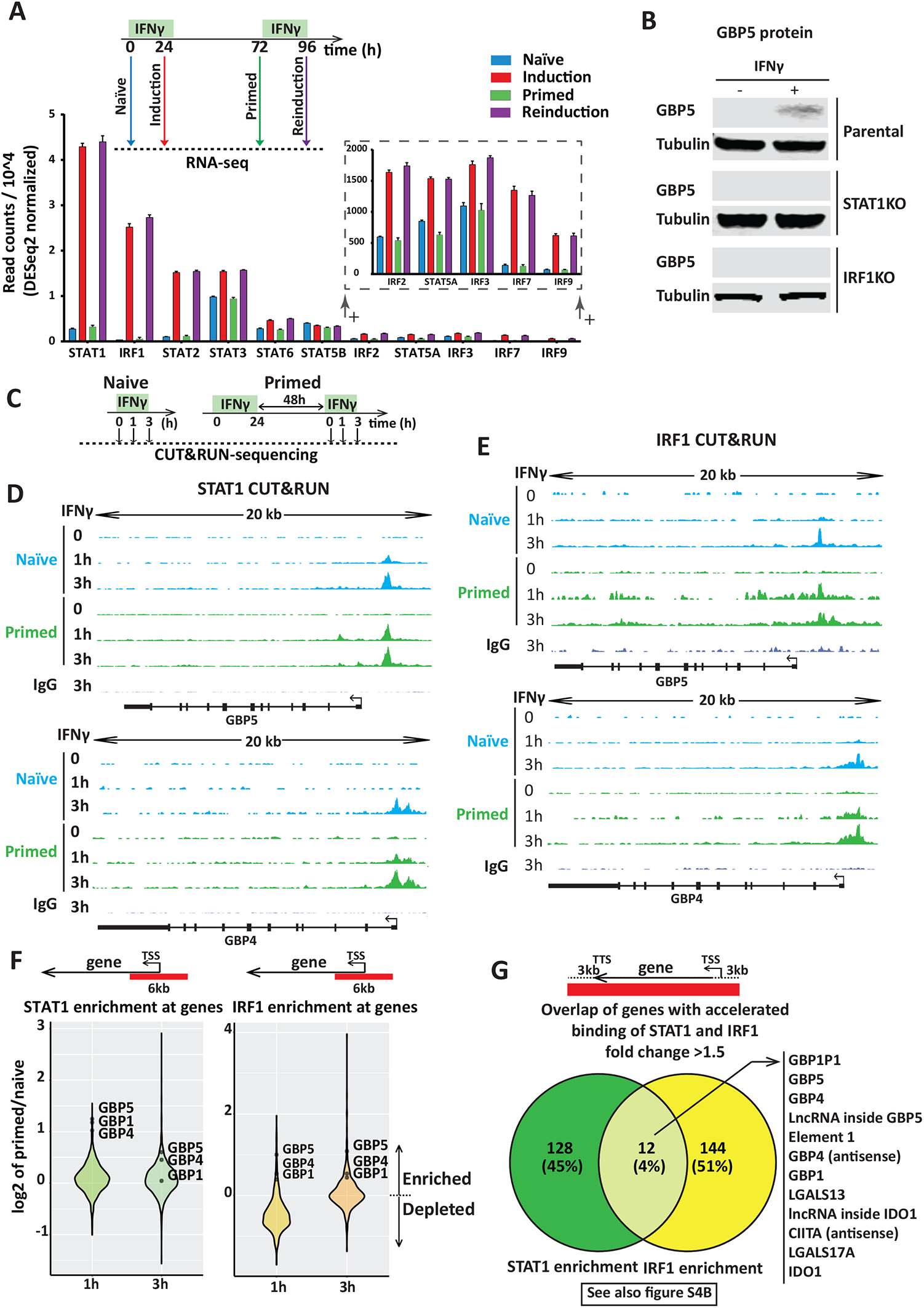
STAT1 and IRF1 are essential for GBP5 expression and show enhanced recruitment to target gene promoters during early reinduction. (A) Top: Experimental outline of IFNγ induction and reinduction regime. Bottom: mRNA levels of STAT and IRF family members at indicated time points based on experiment outlined in the top [obtained from dataset reported in (Siwek et al. 2020)]. Transcription factors are ordered by their expression level. Data are shown as mean. Error bars, SD; n= 3 (**B**) Stable CRISPR knockouts were generated for indicated genes in HeLa cells. Knockout (KO) cells or their parental controls (WT) were induced with IFNγ for 24 hours or left untreated and prepared for SDS-PAGE and immunoblotting. Blots incubated with GBP5 antibody assess gene expression. α-Tubulin (Tubulin) was used as a loading control. **(C)** Scheme describing STAT1 and IRF1 transcription factor enrichment by CUT&RUN **(D-G)** HeLa cells were primed with IFNγ for 24 hours, followed with IFNγ washout. After 48h, naïve and primed cells were induced by IFNγ for 1h and 3h. Cells were harvested at indicated time points and processed for CUT&RUN. Representation of processed data of CUT&RUN for STAT1 **(D)** and IRF1 **(E)** occupancy at GBP5 and GBP4 genes. Sequenced reads were mapped to the human genome (hg38), coverage data are displayed as reads per million (RPM) at equal scaling. **(F)** Violin plot showing log2 fold change of STAT1 and IRF1 enrichment upon treatment of primed relative to naïve cells at the promoter (−3kb to +3 kb relative to TSS) of all annotated genes as measured by CUT&RUN. Data (primed/naïve) is plotted for 1h and 3h IFNγ treatment. **(G)** Venn diagram displaying overlap of STAT1 and IRF1 enrichment for genes (−3kb of TSS and +3kb of TTS) that have more than 1.5-fold change differences between primed and naïve upon 1 hour of IFNγ treatment.

### STAT1 and IRF1 enrichment within the GBP cluster is accelerated during early reinduction

The enhanced promoter accessibility of primed genes upon IFNγ reinduction (Figure S2B) may be driven by a different rate of promoter binding by the essential transcription factors for GBP5 induction, STAT1 and IRF1. To assess this, HeLa cells were induced with IFNγ for 0h, 1h and 3h, both in naïve and primed cells, followed by CUT&RUN (Cleavage Under Targets and Release Using Nuclease) (Meers et al. 2019) for STAT1 and IRF1 (Figure 3C). These results show that STAT1 and IRF1 target the GBP gene promoters within the first three hours of IFNγ induction. Interestingly, both bind faster upon reinduction to primed genes GBP5 and GBP4, GBP1P1 as well as elements distal to GBP5 (Figure 3D, E, S4A). In contrast, both STAT1 and IRF1 target the GBP1 promoter rapidly within an hour of IFNγ that is not further enhanced in primed cells (Figure S4A), consistent with the enhanced chromatin accessibility (Figure S2B). Moreover, unbiased genome-wide analysis in primed vs naïve cells at the promoter of genes (−3kb to +3 kb relative to TSS) revealed that GBP5, GBP1 and GPB4 were among the top loci for accelerated STAT1 and IRF1 recruitment at least at the early time point (Figure 3F).

To explore non-promotor binding sites, we expanded our search to whole genes with 3kb upstream and downstream of the gene body. Interestingly, there are only 12 sites, genome-wide that show accelerated binding of both STAT1 and IRF1 (Figure 3G, S4B), and among those, seven map within the GBP cluster (numbered 1-7 in Figure S4B). Moreover, visual inspection of the GBP cluster identified additional one site with faster STAT1 and IRF1 recruitment in primed cells, which were outside of our defined search range (+/-3kb of gene body), and mapped distal to the GBP5 promoter (Figure 4A, Element 2). Together, these results suggest that faster recruitment of both STAT1 and IRF1 upon reinduction is a feature strongly associated with the in GBP cluster (Figure 4B), both at primed gene promoters, as well as 2 elements 16 kb and 37 kb upstream of the GBP5 promoter. Next, we aimed to determine whether the accelerated binding of both STAT1 and IRF1 in primed cells is the consequence of changes in upstream signalling.

**Figure 4.**
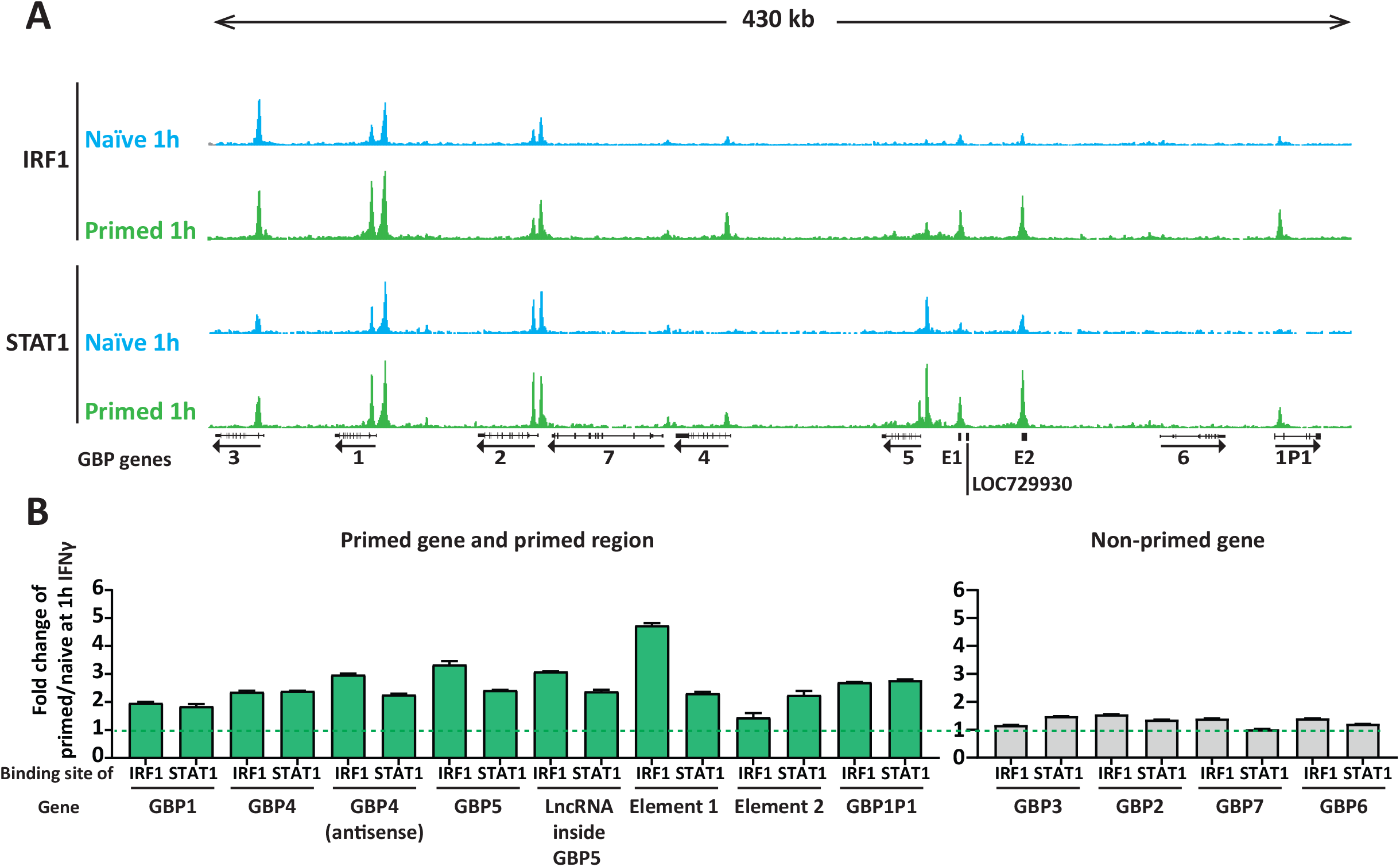
Accelerated recruitment of STAT1 and IRF1 across the primed GBP cluster (A) Tracks of processed CUT&RUN data for STAT1 and IRF1 occupancy across the GBP cluster following 1h of IFNγ induction in naïve and primed cells. Results of sequenced reads were mapped to the human genome (hg38), coverage data are displayed as reads per million (RPM) at equal scaling. GBP gene positions are indicated **(B)** Quantification of STAT1 and IRF1 enrichment in primed relative to naïve cells, 1h after IFNγ induction. Fold change is shown for the 7 loci within the GBP cluster listed in Figure 3G and S4B as well as an additional site (E2) distal to GBP5 (green bars). Enrichment ratios for non-primed genes (grey) are shown for comparison.

### GBP5 priming is not dependent on the regulation of IFNγ-induced STAT1 expression

One possible explanation for GBP5 priming and enhanced promoter binding by STAT1 is STAT1 priming itself. Our RNA-seq analysis shows that STAT1 is strongly induced by IFNγ [(Siwek et al. 2020) and Figure 3A], as previously reported (Cheon and Stark 2009). Although this bulk RNA-seq analysis does not show significant enhancement of expression of STAT1 in primed cells, our single-cell RNA sequencing data from the same study (Siwek et al. 2020), did show a modest degree of priming of STAT1 mRNAs (Figure 5A). To determine whether this effect is relevant for GBP5 priming, we expressed STAT1 from a constitutive promoter in cells in which endogenous STAT1 was deleted (Figure 5B). Effectively, in these cells, STAT1 expression is uncoupled from IFNγ induction and maintained at a level similar to that of induced cells (Figure 5C). Interestingly, while STAT1 is no longer IFNγ-regulated we find that GBP5 expression remains strictly IFNγ-dependent and priming still occurs, indicating that STAT1 expression is not rate-limiting in priming (Figure 5D).

**Figure 5.**
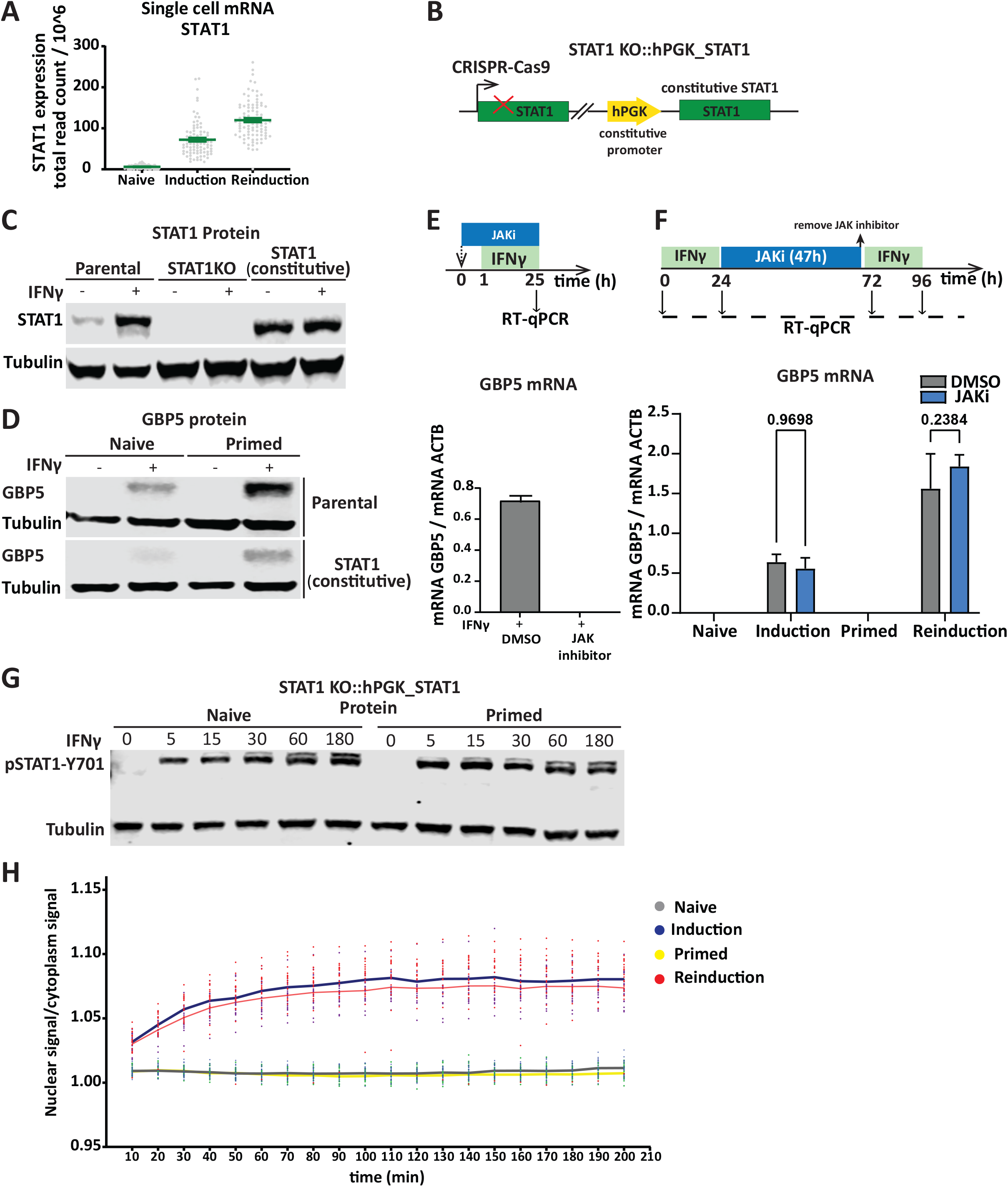
STAT1 expression, activation and import are not rate limiting for priming. (A) Single-cell RNA-seq from HeLa cells for STAT1 from data described in (Siwek et al 2020). Each dot represents STAT1 expression in one cell in naïve (n=91), induction (n=90) and reinduction (n=92) stages. Error bars, SEM. **(B)** Scheme outlining genotype of STAT1 knockout cell line, rescued with constitutive expression of STAT1 from a lentiviral vector. **(C)** Blot probing for STAT1 before and after IFNγ induction in STAT1 knockout (STAT1KO), STAT1 rescued (STAT1, constitutive) and parental control (WT), to confirm knockout and rescue status. α-Tubulin (Tubulin) was used as a loading control. **(D)** STAT1 rescue cells and their parental control were subjected to IFNγ induction and reinduction regime as outlined in Figure 1A. Cell extracts were processed for western blotting and probed for GBP5 expression before and after induction and reinduction as indicated in Figure 1A. α-Tubulin (Tubulin) was used as a loading control. **(E)** Top: Scheme describing HeLa cells treated with JAK inhibitor CP-690550 (JAKi,10 µM) or DMSO vehicle control for 25h together with IFNγ induction for 24h. Bottom: RT-qPCR analysis of GBP5 expression in induced cells treated with JAK inhibitor CP-690550 (JAKi, 10 µM) or DMSO vehicle control. Error bars, SD; n= 3. **(F)** Top: Schematic overview of JAK inhibitor treatment during memory window experiment; HeLa cells primed with IFNγ for 24h, were treated with JAK inhibitor CP-690550 (JAKi, 10 µM) or DMSO vehicle control for 47h, followed by drug washout. After 1h cells were re-induced with IFNγ for 24h. Bottom: GBP5 mRNA level after induction and re-induction in the context of JAKi and DMSO was determined by RT-qPCR and normalized to ACTB mRNA level. Error bars, SD; n= 3. **(G)** Cells constitutively expressing STAT1 (as in B) were primed with IFNγ, followed by IFNγ washout. After 48h, naïve and primed cells were induced by IFNγ for different time points (5, 15, 30, 60, 180 min). Cell extracts were prepared at indicated timepoints and processed for western blotting. Immunoblot of pSTAT1-Y701, and α-Tubulin (Tubulin) as a loading control. **(H)** STAT1-EGFP-dTAG cells were primed with IFNγ, followed by IFNγ washout. After 48h, naïve and primed cells were induced with IFNγ and prepared for live cell imaging. Images were acquired ten minutes after IFNγ addition at 10-minute intervals. The ratio of STAT1-EGFP in nucleus over cytoplasm was quantified. Each dot represents ratio of EGFP fluorescence intensity in one cell. The line shows the mean of data.

### Upstream STAT1 phospho-dynamics and import, are not altered in primed cells

Upon IFNγ stimulation, STAT1 is activated by phosphorylation at Tyr701 via JAK kinase (Darnell, Kerr, and Stark 1994). One possible mechanism of retention of the primed state is that after IFNγ induction, JAK proteins remain active at a low level or stay in a poised state leading to faster activation upon reinduction. To determine the impact of JAK/STAT signalling on GBP gene induction and the maintenance of transcriptional memory, we inhibited JAK kinase using a specific inhibitor (CP-690550, here abbreviated as JAKi). Upon JAK inhibition during IFNγ induction GBP5 expression was lost (Figure 5E), consistent with prior reports (Migita et al. 2011; Ramana et al. 2000). Having established conditions to effectively inhibit JAK, we then treated cells with JAKi immediately following the initial IFNγ induction and kept JAK inhibited during the memory window until just before the second induction (Figure 5F). These results indicate that inhibition of JAK after priming does not significantly affect transcriptional memory indicating that JAK/STAT signalling is required for GBP induction but is dispensable for maintaining memory. While JAK-mediated signalling is not required after priming, it may be hyperactive upon IFNγ exposure, leading to accelerated STAT1 phosphorylation and enrichment on GBP promoters. To determine this, we measured STAT1 Y701 phosphorylation in naïve and primed cells after IFNγ induction at different time points. We find that STAT1 phosphorylation is fast both in naïve and primed cells with a possible minor increase in rate at early timepoints (Figure 5G). To further test if priming has a differential impact on STAT1 activity, we measured STAT1 import into the nucleus which is the downstream consequence of STAT1 activation (Ramana et al. 2002). For this, we employed a cell line constitutively expressing a GFP-tagged STAT1 transgene (described in detail below) and measured STAT1-EGFP levels by live cell imaging (Figure 5H). Treatment of cells with IFNγ leads to STAT1 accumulation in the nucleus, however, STAT1 is not retained in primed cells and nuclear accumulation during induction and reinduction is indistinguishable. We also imaged STAT1 protein by immunofluorescence to quantify nuclear and cytoplasmic pools (Figure S5A). Consistent with the live cell data, the rate of STAT1 import into nucleus is not detectably different between the 1st and 2nd induction and STAT1 is not maintained in the nucleus after removal of IFNγ (Figure S5B). These results indicate that the rates of JAK-mediated STAT1 activation and import are not significantly altered in primed cells compared to naïve cells.

### STAT1 phosphorylation at Ser727 is maintained after removal of IFNγ

In addition to JAK-mediated phosphorylation at Tyr701, STAT1 is also phosphorylated at Ser727 which is required for maximal STAT1 activation and transcriptional activity (Wen, Zhong, and Darnell 1995). A previous study has shown that CDK8, a chromatin-associated kinase known to target Ser727 (Bancerek et al. 2013), remains active in primed cells (D’Urso et al. 2016). We determined the degree of both Tyr701 and Ser727 phosphorylation before and after IFNγ induction in naïve and primed cells. Interestingly, while Tyr701 phosphorylation is rapidly lost after removal of IFNγ, Ser727 phosphorylation is maintained for up to 7 days after priming (Figure 6A, S5C). To test the necessity of STAT1 phosphorylation at Ser727 in transcriptional memory, we rescued the STAT1 knockout cell line (Figure 3B) with a transgene expressing STAT1 in which Ser727 was mutated to alanine (S727A) (Figure S5D). Immunoblotting with phospho-specific antibodies for S727P confirmed loss of phosphorylation (Figure S5E). While STAT^S727A^ expression levels were lower compared to IFNγ-induced levels, this mutant form retained partial functionality as revealed by its ability to support IFNγ-mediated IRF1 induction which is STAT1 dependent (Figure S5E). Nevertheless, this level of STAT1 activity was insufficient to detectably induce GBP5 expression as our readout for transcriptional memory (Figure S5F, G). This indicates that STAT1 phosphorylation at Ser727 plays a critical role in GBP5 expression, which precluded us for further exploring a putative role of Ser727 phosphorylation in IFNγ priming of GBP genes.

**Figure 6.**
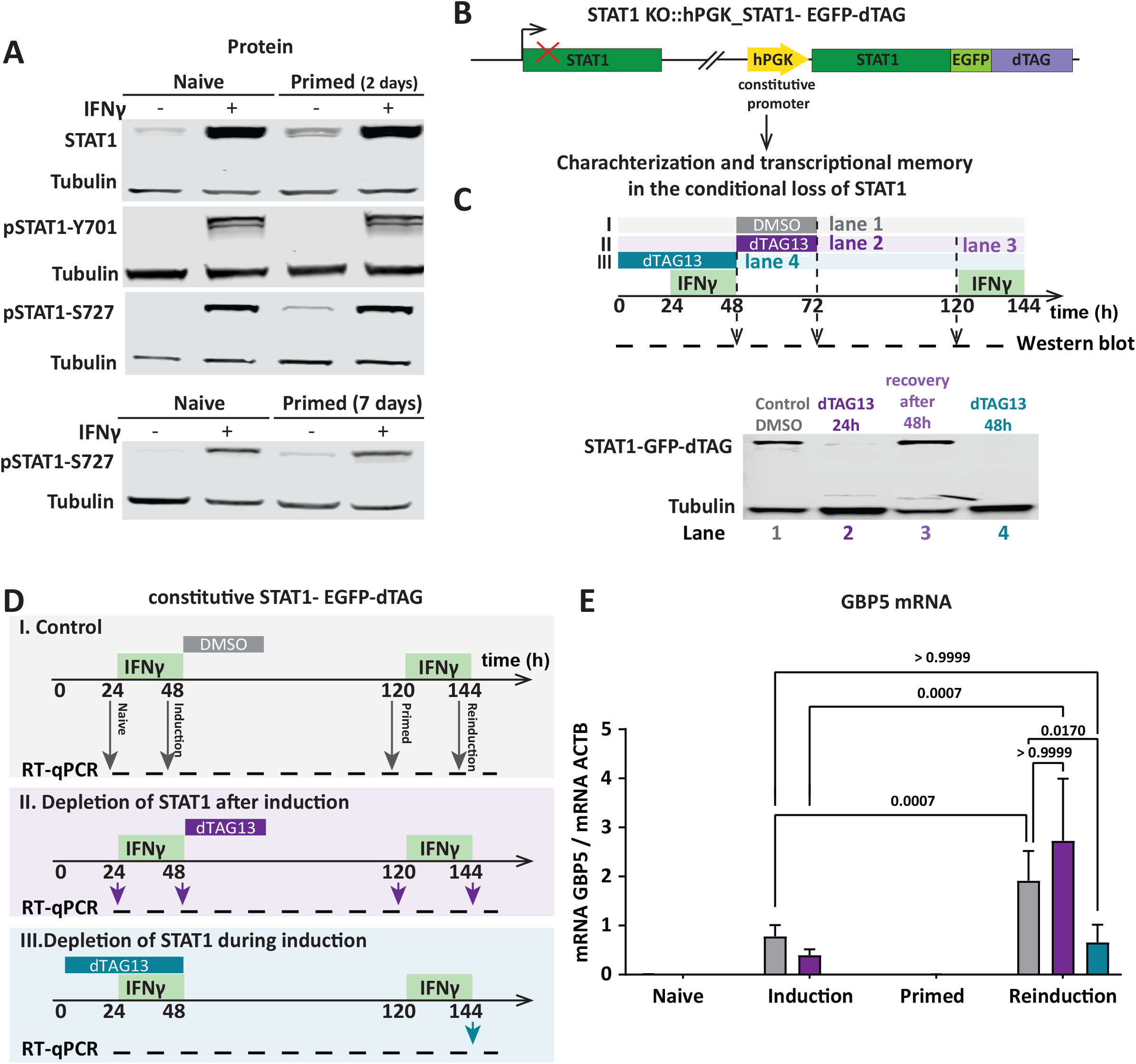
STAT1 phosphorylation at Ser-727 is inherited after removal of IFNγ but not required for priming. (A) HeLa cells were subjected to IFNγ induction and reinduction regime as outlined in Figure 1A with 2 days and 7 days recovery time (primed state) after IFNγ washout. Cell extracts were prepared at indicated timepoints, processed for western blotting and probed for STAT1, pSTAT1-Y701 and pSTAT1-S727. α-Tubulin (Tubulin) as a loading control. **(B)** Schematic overview of STAT1 KO cells rescued with STAT1-EGFP-dTAG (FKB12F36V) under constitutive promoter. **(C)** Characterization of dTAG-induced destruction and recovery. STAT1-EGFP-dTAG expressing cells were induced with IFNγ for 24h, then treated with dTAG13 (100 nM) **(II)** or DMSO vehicle control **(I)** for 24h. Cells were washed three times with medium and after 48h cells were reinduced with IFNγ. In Parallel, **(III)** cells were treated with dTAG13 (100 nM) for 48h and after 24h of drug treatment induced with IFNγ for 24h, then cells were washed three times and after 72h reinduced with IFNγ, cell extracts were processed for western blotting. α-Tubulin (Tubulin) was used as a loading control. **(D)** Experimental outline of dTAG13 depletion and recovery experiment for STAT1 during and after induction [analogous to C but separately outlined for control (I), depletion **after** induction (II) and depletion **during** induction (III)]. **(E)** RNA was isolated, GBP5 mRNA level, as indicated in (D), was determined by RT-qPCR and normalized to ACTB mRNA level. Data are shown as mean (error bars, SD; n=3).

### STAT1 is required during priming to establish GBP5 transcriptional memory but is dispensable during memory of the primed state

Thus far we showed that STAT1 binds to GBP target gene promoters more rapidly in primed cells (Figure 3D) and that STAT1 activation leads to multi-day retention of Ser727 phosphorylation (Figure 6A). However, we could not determine the role of this modification in memory as STAT1^S727^ is critical for GBP5 induction (Figure S5F, G). To circumvent this limitation and directly test the hypothesis that STAT1 itself carries memory of prior IFNγ induction, we constructed a STAT1 allele tagged with a destruction tag (dTAG) composed of a modified FKB12 protein, that can be selectively degraded with the small molecule dTAG13 (Nabet et al. 2018). This allows us to determine whether STAT1 has a role in maintenance of the primed state without affecting its essential role in GBP gene expression upon IFNγ exposure. We expressed STAT1 C-terminally tagged with EGFP-dTAG in HeLa STAT1 knockout cells (Figure 6B). To characterize the functionality of the degron we tested STAT1 dynamics in the context of our IFNγ induction and reinduction regime. Following IFNγ priming, we can effectively remove the vast majority of STAT1 within 24 hours of dTAG13 addition, while DMSO controls retain STAT1 levels (Figure 6C). By 48h post removal of dTAG13, STAT1 levels are fully recovered, prior to IFNγ reinduction. Having established conditions in which we can selectively remove STAT1 only after priming and re-express prior to reinduction, we determined the effect of STAT1 removal on transcriptional memory (Figure 6D). We find that STAT1 depletion immediately after priming does not impair GBP5 memory and allows for enhanced GBP5 expression upon IFNγ reinduction, similar to controls (Figure 6E). These results demonstrate that STAT1 and its associated posttranslational modifications are not necessary for maintenance of GBP5 transcriptional memory. To further explore the role of STAT1, we aimed to determine the requirement of STAT1 during priming. While STAT1 is necessary for GBP5 induction, whether it plays a role in priming has not been determined. Possibly, IFNγ signalling leads to priming via factors other than STAT1. To test this directly, we depleted STAT1 with dTAG13 prior to, and during IFNγ priming (Figure 6D). We then let STAT1 levels recover before IFNγ reinduction. Strikingly, under these conditions, GBP5 expression is not enhanced after the 2^nd^ IFNγ induction. Despite STAT1 presence during reinduction, the cells behave as if naïve despite prior exposure to IFNγ (Figure 6E). These results demonstrate that STAT1 is necessary during cell priming, not only to induce GBP expression but also to establish transcriptional memory.

## Discussion

In this report, we dissected the role of STAT1 in the establishment and maintenance of IFNγ-induced transcriptional memory. We focussed on GBP5, GBP4 and GBP1 as target genes that show strong priming upon IFNγ induction, as previously described (Gialitakis et al. 2010; Siwek et al. 2020) to understand transcriptional memory (Figure 1A). Prior work showed that RNA polymerase II (Pol II) is retained at promoters following IFNγ priming (Light et al. 2013) or is recruited more rapidly following reinduction (Kamada et al. 2018). An increase in both chromatin accessibility (Figure S2) and Pol II recruitment (Kamada et al. 2018) can be due to alteration of local chromatin structure induced by transcription, regardless of upstream signalling. However, we find that transcription per se, induced using CRISPRa-SAM-mediated gene activation is not sufficient to provoke the primed state (Figure 2). Artificial CRISPRa-mediated gene activation is known to bring RNA Pol II, the general transcription machinery and increases the acetylation levels on nucleosomes (Giménez et al. 2016). Interestingly recruitment of these factors is not sufficient to prime the GBP target gene, implying a specific IFNγ-mediated factor is required. Furthermore, these results suggest that while RNA polymerase II has been detected on promoters of primed HLA genes (Light et al. 2013), it may itself not be the primary driver of transcriptional memory. In line with this, recent elegant experiments in yeast have shown that in the case of *INO1* transcriptional memory, RNA polymerase II is poised but not required for memory (Sump et al. 2022).

We showed both STAT1 and IRF1 are required for GBP5 induction, and both display an accelerated recruitment to the promoters of primed genes during reinduction (Figure 3). Intriguingly, the set of loci that show enhanced recruitment of both STAT1 and IRF1 are largely restricted to the GBP cluster (Figure 3G, 4). We excluded priming of STAT1 expression or the presence of STAT1 protein to be involved in the maintenance of memory (Figure 5D, 6E). Combined, these results lead us to propose that the primed state, induced by IFNγ exposure, does not involve upstream JAK-STAT signalling nor is a consequence of mere target gene expression. Instead, memory appears to be restricted to a state induced by STAT1 that may include changes in local chromatin structure, specifically at memorized GBP genes that facilitate accelerated recruitment of STAT1 and IRF1. Interestingly, we detected Ser727 phosphorylation of STAT1 in primed cells (Figure 6A, S5C) for up to 7 days of memory. While our STAT1 depletion experiments demonstrate that this phosphorylation is not the carrier of the primed state (Figure 6E), it is possible that the underlying chromatin maintains the capacity to rapidly active STAT1 upon rebinding to promoters. In this vein, it is noteworthy that the putative kinase for STAT1 Ser727, CDK8 is maintained on target gene chromatin in IFNγ-primed cells (‘D’Urso et al. 2016) and that CDK8 occupancy correlates with STAT1 S727 phosphorylation (Bancerek et al. 2013). Other possible changes in chromatin structure and componsition may include SMARCA4 (BRG1), a SWI/SNF related remodelling complex (Sif et al. 2001) that can be recruited by STAT1 to GBP genes (Ni et al. 2005).

In sum, our work defines that IFNγ priming results in STAT1-dependent changes of primed genes, excluding effects in upstream signalling or downstream transcription activation. This focusses future efforts on identifying what factors cause local chromatin changes that enhance target gene expression upon IFNγ re-exposure.

## Author Contributions

S.S.H.T., W.S. and L.E.T.J. conceived the study. S.S.H.T. designed and performed the experiments, analyzed the data, and created the figures. W.S. designed and performed RNA-seq experiments. P.M. helped to analyze the data for the CUT&RUN experiment. J.F.M and I.A.Z. provided tissue culture and experimental expertise and assisted with cloning. L.E.T.J. and W.S. supervised the work and helped to design the experiments and interpret the data.

L.E.T.J. acquired funding. S.S.H.T. and L.E.T.J wrote the manuscript.

## Conflict of Interest Declaration

The authors declare that they have no conflict of interest.

Acknowledgments

This work was funded by an ERC-consolidator grant ERC-2013-CoG-615638 and a Senior Wellcome Research Fellowship 210645/Z/18/Z to LETJ. SSHT was supported by a Fundação para a Ciência e a Tecnologia (FCT) doctoral fellowship PD/BD/128438/2017. WS was supported by a FCT postdoctoral fellowship (SFRH/BPD/117179/2016) and a Marie Sklodowska-Curie individual fellowship (101025900). LETJ receives salary support from the University of Oxford, Department of Biochemistry and St Edmund Hall.

## Experimental procedures

Resource table:
| REAGENT or RESOURCE | SOURCE | IDENTIFIER |
| --- | --- | --- |
| **Experimental Models: Cell Lines** |  |  |
| Human: HeLa WT | ATCC | Cat#CCL-2; RRID: CVCL_0030 |
| Human: HeLa Kyoto | https://web.expasy.org/cellosaurus/<br>CVCL_1922 | RRID: CVCL_1922 |
| Human: HEK293 | ATCC | CRL-1573; RRID: CVCL_0045 |
| Human: HeLa STAT1 knockout polyclonal | this study | N/A |
| HeLa STAT1 knockout clonal | this study | N/A |
| HeLa constitutive STAT1 polyclonal | this study | N/A |
| HeLa constitutive STAT1<sup>S727A</sup> polyclonal | this study | N/A |
| HeLa constitutive STAT1-GFP-FKB12<sup>F36V</sup> (dTAG13) polyclonal population | this study | N/A |
| HeLa STAT2 knockout polyclonal | this study | N/A |
| HeLa STAT3 knockout polyclonal | this study | N/A |
| HeLa STAT5B knockout polyclonal | this study | N/A |
| HeLa IRF1 knockout polyclonal | this study | N/A |
| HeLa IRF9 knockout polyclonal | this study | N/A |
| **Recombinant DNA** |  |  |
| lentiCRISPR v2 | Sanjana et al., 2014; Addgene | Addgene #52961 |
| psPAX2 | Addgene | Addgene #12260 |
| pMD2.G | Addgene | Addgene #12259 |
| lenti sgRNA(MS2)_Puro | Addgene | Addgene #73795 |
| lentiMS2-P65-HSF1_Hygro | Addgene | Addgene #61426 |
| lenti dCas9-VP64_Blast | Addgene | Addgene #61425 |
| pRosetta | Addgene | Addgene #59700 |
| pRosetta_STAT1 | This study | N/A |
| pRosetta_STAT1<sup>S727A</sup> | This study | N/A |
| EGFP-dtag-HA | Jansen Lab | N/A |
| **Antibodies** |  |  |
| GBP5 | Cell Signaling Technology | Cat#67798 |
| GBP5 | Abcam | Cat#ab96119 |
| EGFP | Merck | Cat#G1544 |
| α-TUB | Merck | Cat#T9026 |
| Anti-rabbit (fluorophore conjugated) | LI-COR | Cat#926-32211 |
| Anti-mouse (fluorophore conjugated) | Rockland | Cat#610-744-124 |
| STAT1 | Cell Signaling Technology | Cat#9172 |
| pSTAT1 (701) | Cell Signaling Technology | Cat#9167 |
| pSTAT1 (727) | Cell Signaling Technology | Cat#9177 |
| STAT2 | Santa Cruz Biotechnology | Cat#sc-1668 |
| STAT3 | Santa Cruz Biotechnology | Cat#sc-8019 |
| STAT5B | Santa Cruz Biotechnology | Cat#sc-1656 |
| IRF1 | Cell Signaling Technology | Cat#8478 |
| IRF9 | ThermoFisher Scientific | Cat#702322 |
| **Oligonucleotides and other sequence-based reagents** |  |  |
| CRISPRa gRNAs | This study | Table 1 |
| CRISPR/Cas9 gRNAs | This study | Table 1 |
| qPCR primers | This study | Table 1 |
| GeneArt Strings DNA Fragments (STAT1 cDNA) | ThermoFisher | Supplementary 6 |
| **Bacterial and Virus Strains** |  |  |
| ER2566 | Jansen Lab | N/A |
| DH5alpha | Jansen Lab | N/A |
| **Chemicals, Peptides, and Recombinant Proteins** |  |  |
| Interferon gamma (IFNγ) | Merck | Cat#SRP3058; CAS: 9008-11-1 |
| Vybrant DyeCycle Ruby Stain | ThermoFisher | Cat # V10309 |
| Tofacitinib citrate | Merck | Cat# PZ0017 |
| dTAG13 | Merck | Cat #SML2601-1MG |
| T4 DNA Polymerase | New England Biolabs | Cat#M0203S |
| Lipofectamine LTX with Plus Reagent | ThermoFisher | Cat# A12621 |
| **Software** |  |  |
| CellProfiler v. 3.1.9 | https://cellprofiler.org/ | N/A |
| ImageJ v. 1.52h | https://imagej.nih.gov/ij/ | N/A |
| GraphPad Prism (no version?) | https://www.graphpad.com/scientific-software/prism/ | N/A |
| RStudio v. 1.3.1093 | https://www.rstudio.com/ | N/A |
| Deeptools v. 3.5.0 | https://anaconda.org/bioconda/deeptools | N/A |
| Samtools v. 1.15 | https://anaconda.org/bioconda/samtools | N/A |
| Bowtie2 v. 2.4.5 | https://anaconda.org/bioconda/bowtie2 | N/A |
| GPP sgRNA Designer software | https://portals.broadinstitute.org/gpp/ | N/A |
| Biorender | https://biorender.com/ | N/A |
| **Other (Kits, instrumentation, laboratory equipment, lab ware etc)** |  |  |
| Monarch® RNA Cleanup Kit | New England Biolabs | Cat#T2040L |
| RNA-to-cDNA Kit | Applied Biosystems | Cat#4387406 |
| CUTANA™CUT&RUN kits | EpiCypher | Cat#14_1048 |
| NEBNext Ultra II DNA Library Prep Kit for Illumina | New England Biolabs | Cat#E7645S |
| TDE1 Tagment DNA Enzyme | Illumina | Cat#15027865 |

### Resource table

#### Human Cell Lines

- HeLa (female, RRID: CVCL_0030); used in Figure 1, 2, 3, 5, 6, S2, S3, S5
- HeLa Kyoto (female, RRID: CVCL_1922); used in Figure 2, 3, 4, S1, S4
- HeLa STAT1 knockout polyclonal; used in Figure 4
- HeLa STAT1 knockout clonal; used in Figure 5, S5E
- HeLa constitutive STAT1 polyclonal; used in Figure 5
- HeLa constitutive STAT1 ^S727A^ polyclonal; used in Figure S5
- HeLa constitutive STAT1-GFP-FKB12 F36V (dTAG13) polyclonal; used in Figure 5, 6
- HeLa STAT2 knockout polyclonal; used in Figure S3
- HeLa STAT3 knockout polyclonal; used in Figure S3
- HeLa STAT5B knockout polyclonal; used in Figure S3
- HeLa IRF1 knockout polyclonal; used in Figure 3, S3
- HeLa IRF9 knockout polyclonal; used in Figure S3
- HEK239T (female, RRID: CVCL_0045); used for virus production

Cell lines and culture conditions

All cell lines were incubated at 37°C, 5% CO_2_, and grown in ‘Dulbecco’s Modified Eagle Medium (DMEM) containing high glucose and pyruvate (ThermoFisher, 41966-029) supplemented with 10% NCS (newborn calf serum, ThermoFisher, 16010-159) and 1% Penicillin-Streptomycin (ThermoFisher, 15140-122). For passaging, cells were washed using 1X DPBS (ThermoFisher), detached using TrypLE Express phenol red (ThermoFisher) and resuspended in DMEM. Cells were counted using Countess™ Cell Counting according to the manufacturer’s instructions (ThermoFisher Scientific). Transfection of cells was performed using Lipofectamine LTX (ThermoFisher Scientific) according to the manufacturer’s instructions.

### Method details Reagents

All chemicals, unless otherwise noted, were obtained from ThermoFisher or Merck. Enzymes were obtained from New England Biolabs. The following drugs/dyes were used for this work: IFNγ (final concentration of 50ng/mL, Merck), dTAG13 (final concentration of 100nM, Merck), Vybrant DyeCycle Ruby Stain (final concentration of 5µM, ThermoFisher) and Tofacitinib citrate also known as CP-690550 (JAK inhibitor, concentration of 10µM, Merck). The following antibodies were used for this work: GBP5 (Cell Signaling Technology, 67798; Abcam, ab96119), STAT1 (Cell Signaling Technology, 9172), Phospho-STAT1 (Ser727) (Cell Signaling Technology, 9177), Phospho-STAT1 (Tyr701) (58D6) (Cell Signaling Technology, 9167), STAT2 (Santa Cruz Biotechnology, sc-1668), STAT3 (Santa Cruz Biotechnology, sc-8019), STAT5B (Santa Cruz Biotechnology, sc-1656), IRF1 (Cell Signaling Technology, 8478), IRF9 (ThermoFisher Scientific,702322), a-TUB (Merck, T9026), anti-rabbit (fluorophore conjugated) (LI-COR, 926-32211), anti-mouse (fluorophore conjugated) (Rockland, 610-744-124).

### DNA constructs and genome engineering

A constitutively expressed STAT1 was constructed in the pRosetta plasmid (Addgene ##59700). First, the EGFP sequence was deleted by PCR from the plasmid backbone using primers (5’
s-gaagcggagctactaacttcagcctgctgaagcagg-3’, 5’-ggtggatccccctggggagagaggtcg-3’). The product was blunted by T4 DNA Polymerase (New England Biolabs) and self-ligated with T4 DNA Ligase (New England Biolabs). A synthetized fragment was cloned into this vector carrying the cDNA of the α isoform of STAT1 with a silent PAM site mutation (GeneArt Strings, sequences in supplementary 6) and 25 bp homology arms by the SLIC method (Jeong et al. 2012).

To construct the STAT1^S727A^ variant in the pRosetta-STAT1 plasmid, mutations were introduced using the primers 5’
s-ACAACCTGCTCCCCATGGCTCCTGAGGAGTTTGACG-3’ and 5’-CGTCAAACTCCTCAGGAGCCATGGGGAGCAGGTTGT-3’, replacing Serine 27 in STAT1 to Alanine. (The underline nucleotides are the introduced mutations)

To construct the STAT1-GFP-FKB12 F36V (dTAG13), the EGFP-FKB12 F36V fragment was first amplified from EGFP-dtag-HA plasmid using primers (5’
s-GAATTCGACAGTATGATGAACACAGTAGCCATGGTGAGCAAGGGCGAGGAG-3’, 5’-AGGCTGAAGTTAGTAGCTCCGCTTCCGCTAGGTGCATAGTCCGGGACATCATACG -3’) and inserted into pRosetta-STAT1 plasmid in frame with the STAT1 C-terminus by the SLIC method (Jeong et al. 2012).

All plasmid inserts were verified by DNA sequencing. Expression plasmids were co-transfected with lentiviral packaging plasmid psPAX2 (Addgene #12260), and envelope plasmid pMD2.G (Addgene #12259) into HEK293T cells at a molar ratio of 4:3:1, respectively. Lentiviral particles were harvested as described (Dull et al. 1998). The lentivirus containing the desired construct were then transduced into HeLa cells (see below in CRISPR/Cas9 knock-out and lentivirus packaging section).

### Transcriptional memory assay

Cells were primed with IFNγ (Merck) or left untreated for 24h, followed by IFNγ washout with DPBS (ThermoFisher) and trypsinization by TrypLE (ThermoFisher) to harvest cells. Cells were cultured with fresh medium for another 48h, unless stated otherwise. Next, naïve and primed cells were induced by IFNγ for 24h. After 24h, cells were trypsinized and harvested, and the pellets were processed for subsequent experiments.

### ATAC-seq

The ATAC-seq procedure was adapted from Omni-ATAC protocol (Corces et al. 2017). Briefly, 50,000 naive or primed cells treated with IFNγ for 1h and 3h and non-treated cells were centrifuged for 5 minutes at 500*g*, resuspended in 50μL of cold ATAC lysis buffer (10mM Tris-HCl pH 7.4; 10mM NaCl; 3mM MgCl_2_; 0.1% NP-40; 0.1% Tween-20 and 0.01% Digitonin), and incubated on ice for 3 minutes. Next, 1mL wash buffer was added to the pellets (10mM Tris-HCl pH 7.4; 10mM NaCl; 3mM MgCl2; 0.1% Tween-20), cells were resuspended and immediately centrifuged for 10 minutes at 500*g* at 4°C. Next cell pellets were resuspended in 50μL tagmentation reaction buffer [25μL 2 × TD buffer, 16.5μL 1X PBS, 0.1% Tween-20, 0.01% Digitonin, 5.4μL nuclease-free water and 2.5μL TDE1 (Tn5 enzyme, Illumina)] followed by incubation at 37°C for 30 minutes. DNA was then purified using MinElute PCR Purification Kit (QIAGEN) (10μL elution).

DNA library preparation was performed as previously reported (Siwek et al. 2020). In brief, the purified DNA was amplified by Q5 Hot start DNA polymerase using described indexing primers (Buenrostro et al. 2013). The thermal cycling process was programmed as follows:n °C for 5 minutes and 98°C for 30s, and five cycles of (98°C 10s; 63°C 30s; 72°C 1 minute); followed by subsequent library quantification using qPCR with the following program: 95°C for 30s, 95°C for 10s, 58°C for 30s, 72°C for 1 minute, and additional PCR amplification that is needed for each sample to reach 1/3 of saturated signal. DNA library was purified and size selected using a double-sided bead purification protocol with AMPure XP beads that removes both unwanted small and large fragments (Beckman Coulter). The QuBit dsDNA HS Assay (ThermoFisher Scientific) was used to determine DNA concentration according to the manufacturer’s protocol. Fragment size was estimated by DNA TapeStation (Agilent High Sensitivity D1000) prior to mixing of multiplexed libraries diluted to 2nM concentration (calculated based the on Qubit dsDNA HS Assay Kit) for sequencing using the NextSeq 500/550 v2.5 kit (Illumina, 75bp single-end reads). Finally, sequenced reads were mapped to the human genome (hg38) using bowtie2 (Langmead and Salzberg, 2012). Coverage bigwig files were generated using bamCompare (Ramírez et al., 2016), with 50bp bin size. Data was normalized to reads per million (RPM).

### CRISPRa-SAM

CRISPRa-SAM transfection was performed as described (Konermann et al. 2014) with the following modifications: Guide RNAs (gRNAs) for GBP1 and GBP5 were designed using GPP Web Portal (Broad Institute). gRNAs were designed 200-300 bases upstream of the TSS. All oligo sequences can be found in Supplementary Table 1. Cells were transiently transfected with Cas9 component plasmid (Addgene #61425), gRNAs plasmid (Addgene #73795) and MS2-P65-HSF1 activator plasmid (Addgene #61426) at a molar ratio of 2.5:1:10, respectively. Lipofectamine LTX (ThermoFisher) transfection was performed according to the manufacturer’s protocol. After 4h, the cells were cultured in fresh medium and incubated for 2 days at 37°C. Transfected cells were harvested and processed for further experiments.

### CRISPR/Cas9 knock-out and lentivirus packaging and transfection

To mutate STAT and IRF genes, gRNAs were selected from the Toronto KnockOut (TKO) CRISPR Library (Supplementary Figure 6). The lentiCRISPR plasmids (Addgene #52961) with cloned gRNAs were co-transfected (as described above) with lentiviral packaging plasmid psPAX2 (Addgene #12260) and envelope plasmid pMD2.G (Addgene #12259) into HEK293T cells at a molar ratio of 4:3:1, respectively. Cells were incubated for 3 days at 37°C, prior to collecting the medium containing the virus and filtering through a 0.45μm filter. HeLa cells were incubated in medium containing 8μg/mL of polybrene (Merck) for 1h, then infected with filtered viruses carrying Cas9 with gRNAs targeting STAT and IRF genes. The cells were left to grow for 48h followed by selection with puromycin (1μg/mL).

### RT-qPCR

RNA was extracted using TRIzol (Invitrogen) based on manufacturer’s instructions. DNA was removed with DNase I (New England Biolabs) and RNA was purified using Monarch RNA Cleanup Kit (New England Biolabs). Complementary DNA (cDNA) was generated using High-Capacity RNA-to-cDNA Kit (Applied Biosystems) and the cDNAs were diluted 10-fold prior to qPCR measurements. The qPCR assay was performed with iTaq Universal SYBR Green Supermix (BioRad) on CFX384 Real-Time System machine (BioRad). qPCR primers are listed in Supplementary Table 1. All experiments were performed in technical and biological triplicates. Primer efficiency and qPCR quantification were analyzed as previously described (Siwek et al. 2020).

### Immunoblotting

Cell pellets were resuspended in protein sample buffer (125mM Tris-HCl pH 6.8, 10% Glycerol, 1% SDS, 0.2% (w/v) Orange G, 5% β-mercaptoethanol) and incubated in 98°C for 5 minutes. Benzonase (50U) was added to the lysates and incubated at room temperature for 30 minutes, followed by incubation in 98°C for 5 minutes and 10 minutes centrifugation. Soluble extracts were separated on a 10% or 12% SDS-PAGE gel (Bio-Rad), then transferred to nitrocellulose membranes (BioRad Transblot Turbo), blocked with Intercept (PBS) Blocking Buffers (LI-COR) for 1h and incubated overnight with primary antibodies at 4°C. The following day, blots were washed three times with TBST (20mM Tris-HCl pH 7.5, 150mM NaCl, 0.1% Tween 20) and incubated with secondary antibodies for 1h, with subsequent three times TBST washing. The blots were analyzed by Odyssey Imaging System (LI-COR) and quantified using ImageStudio.

### CUT&RUN

For CUT&RUN (Meers, Bryson, and Henikoff, 2019), samples were processed using CUTANA™ CUT&RUN kit (EpiCypher) according to the manufacturer’s instructions. In brief, cells were fixed with 1% formaldehyde for 1 minute at room temperature, followed by quenching in 25mL of 125mM glycine in PBS. Cells were harvested by low-speed centrifugation, 500,000 cells were washed with washing buffer, then incubated with pre-activated Concanavalin A-coated beads for 10 minutes at room temperature, followed by an overnight incubation at 4°C with antibodies (0.5μg) in antibody-buffer containing 0.01% digitonin. Next, ConA beads bound cells were washed twice with permeabilization buffer containing 0.01% digitonin, then incubated with protein A-MNase fusion for 10 minutes at room temperature and washed to remove unbound protein A-MNase. Cleavage was performed at 4°C for 2h by addition of calcium chloride to a final concentration of 100mM. After incubation, reaction was stopped by addition of the STOP buffer (containing fragmented genomic *E. coli* DNA as spike-in). Fragmented DNA samples were extracted after 10 minutes of incubation at 37°C, followed by phenol-chloroform extraction. For preparation of DNA libraries, fragmented DNA was processed using NEBNext Ultra II DNA Library Prep Kit for Illumina (New England Biolabs) according to the manufacturer’s instructions. Next, the material was purified and size selected with AMPure XP beads (double-sided bead purification protocol, Beckman Coulter). Purified multiplexed libraries were diluted to 2nM concentration (calculated based on the Qubit dsDNA HS Assay Kit) for sequencing using NextSeq 500/550 v2.5 Kit (Illumina, 35bp paired-end reads). Finally, sequenced reads were mapped to the human genome (hg38) using bowtie2 (Langmead and Salzberg, 2012). Coverage bigwig files were generated using bamCompare (Ramírez et al., 2016), with 50bp bin size. Coverage data was normalized to reads per million (RPM).

### Immunofluorescence

Immunofluorescence protocols were adopted from (Bodor et al. 2012). In brief, cells were fixed on poly-l-lysine coated glass coverslips with 4% formaldehyde (ThermoFisher Scientific) for 10 min followed by permeabilization with in PBS with 0.1% v/v Triton-X-100 (PBS-TX) (ThermoFisher), blocked for 30 minutes at 37°C, then incubated with STAT1 antibody (1:50, Cell Signaling Technology) for 1h at 37°C. Coverslips were washed with PBS-TX and incubated with fluorescein conjugated anti-rabbit IgG (1:200, Rockland Immunochemicals) for 30 min at 37°C. Nuclei were stained using DAPI (Merck). Coverslips were mounted in Mowiol and stored at 4°C until imaging.

### Microcopy

Imaging was performed on Leica Microsystems DMI 6000B inverted-light microscope at 40X magnification using a 1.4 NA oil objective (HC PLAN APO) to capture 0.2μm Z-stacks. Images were quantified using ImageJ macros. CRaQ is described previously (Bodor et al. 2012), and modified to calculate the whole DAPI signal as region of interests (ROIs). The macro quantifies median of whole cell STAT1 levels and nuclear STAT1 levels (using DAPI as mask).

Live cell imaging was performed on Leica Microsystems DMI 6000B inverted-light microscope at 20X magnification, a microscope stage incubator maintained at 37°C in a humidified atmosphere. HeLa STAT1 KO cells constitutively expressing STAT1-GFP-FKB12 F36V (dTAG13) were seeded in chamber slides (ThermoFisher) and grown in Live Cell Imaging solution (ThermoFisher) containing 10% FBS and 1% Penicillin-Streptomycin (ThermoFisher, 15140-122). Vybrant DyeCycle Ruby Stain (ThermoFisher) was added 1h before imaging at a final concentration of 5 µM to mark cell nuclei. Images were acquired every 10 minutes immediately after IFNγ induction and quantified using CellProfiler (Carpenter et al. 2006).

### FACS

Cells were harvested and re-suspended in ice-cold conditional medium (1:1 mixture of fresh complete medium and filtered medium collected from proliferating cell cultures) supplemented with 20% Fetal Bovine Serum, 0.25mg/mL Fungizone (ThermoFisher Scientific), 1% Penicillin-Streptomycin (ThermoFisher, 15140-122)), and filtered through a 5mL polystyrene round-bottom tubes with cell-strainer caps (Falcon) before sorting (FACSCalibur) (BD Biosciences). Cells were collected in conditional medium.

### Quantification and statistical analysis

RT-qPCR and STAT1 CUT&RUN data were collected in triplicate, IRF1 CUT&RUN data were collected in duplicate (with the exception of the IRF1 primed 3h data point for which only one replicate is shown). For ATAC-seq one replicate experiment is shown. Standard deviation is reported. Statistical analyses and P value calculations for the RT-qPCR data were performed by one-way analysis of variance (ANOVA) (GraphPad prism, v9.3.1) and can be found in the figures.

## Figure legends

**Supplementary Figure 1. Titration of minimal IFNγ pulse to prime GBP1. (A)** Experimental outline of GBP1 priming with different concentrations of IFNγ and incubation times. HeLa cells were induced with 1, 3 or 50 ng/mL of IFNγ for 4h and 24h, followed by IFNγ washout. After 48h, naïve and primed cells were induced with IFNγ (50ng/mL) for 24h and harvested for GBP1 mRNA analysis by RT-qPCR after induction **(B)** and reinduction **(C)**, normalized to ACTB mRNA level.

**Supplementary Figure 2. Promoter accessibility of GBP genes upon IFNγ stimulation. (A)** Scheme describing chromatin accessibility (ATAC-seq) experiment. HeLa cells were primed with IFNγ for 24 hours, followed by IFNγ washout. After 48h, naïve and primed cells were induced by IFNγ for 1h and 3h. Cells were harvested at indicated time points and processed for ATAC-seq. **(B)** Results of sequenced reads were mapped to the human genome (hg38), coverage data is displayed as reads per million (RPM) at equal scaling for two genes showing priming, GBP1, GBP4 and GBP5 **(Left)**, two IFNγ-induced genes, IRF1 and TAP1 **(Right)**. The data across samples are scaled equally for each locus. **(C)** No nearby enhancers are accessible in primed cells. Representation of processed data for ATAC-seq at the GBP5 gene and upstream region. Sequenced reads were mapped to the human genome (hg38), coverage data are displayed as reads per million (RPM) at equal scaling.

**Supplementary Figure 3. Knock out cell lines of all relevant STAT and IRF genes to determine requirement for GBP5 expression**. Stable CRISPR knockouts were generated for indicated genes in HeLa cells. Knockout (KO) cells and their parental controls (WT) were induced with IFNγ for 24h or left untreated. **(A)** Immunoblots probing for STAT and IRF transcription factors to confirm knockout status, **(B)** probing for the effect on GBP5 expression **(C)** probing the effect on GBP5 priming as outlined in Figure 1A. α-Tubulin (Tubulin) was used as a loading control.

**Supplementary Figure 4. Accelerated STAT1 and IRF1 binding in primed genes (A)** Representation of processed CUT&RUN data for STAT1 and IRF1 occupancy at the selected GBP genes loci (GBP1P1, LOC729930 and GBP1). **(B)** List of loci showing enhanced STAT1 and IRF1 binding in primed cells as in Figure 3G (−3kb of TSS and +3kb of TTS with a minimum of 1.5-fold difference between primed and naïve upon 1 hour of IFNγ treatment).

**Supplementary Figure 5. Priming does not change the rate of STAT1 import and phosphorylation at Ser727 which essential for GBP5 expression. (A)** Constitutive STAT1 expressing cells were subjected to IFNγ induction and reinduction regime as outlined in Figure 1A. Cells were fixed following indicated treatments as in Figure 1A, followed by immunostaining for STAT1 and DAPI. **(B)** Quantification of ratio of STAT1 in nucleus over cytoplasm in fixed cells. Data are shown as mean (error bars, SD; n=10) **(C)** Replicates of experiments shown in Fig. 6A, blots probed for pSTAT1-S727 and α-Tubulin (Tubulin) as a loading control. **(D)** Schematic overview of STAT1 KO cell line rescued with a STAT1 variant with a S727A mutation under constitutive promoter. **(E)** STAT1KO::STAT1-S727A expressing cells or their parental controls (WT) were induced with IFNγ for 24h or left untreated and processed for western blotting. Extracts were probed for STAT1, pSTAT1-S727 and IRF1 to confirm the mutation and to assess STAT1-S727A function. **(F)** STAT1-S727A expressing cells or their parental controls (WT) were subjected to IFNγ induction and reinduction regime as outlined in Figure 1A, cell extracts were immunoblotted to determine the GBP5 protein level. α-Tubulin (Tubulin) was used as a loading control. **(G)** in parallel to (F), RNA was isolated and GBP5 mRNA level was determined by RT-qPCR and normalized to ACTB mRNA level. Data are shown as mean (error bars, SD; n=3).

**S6. Table 1. List of gRNAs and primes.**

## References

Antonczyk, Aleksandra, Bart Krist, Malgorzata Sajek, Agata Michalska, Anna Piaszyk-Borychowska, Martyna Plens-Galaska, Joanna Wesoly, and Hans A.R. Bluyssen. 2019. “Direct Inhibition of IRF-Dependent Transcriptional Regulatory Mechanisms Associated with Disease.” Frontiers in Immunology 10 (MAY): 1176. https://doi.org/10.3389/FIMMU.2019.01176/BIBTEX.

Bancerek, Joanna, Zachary C. Poss, Iris Steinparzer, Vitaly Sedlyarov, Thaddäus Pfaffenwimmer, Ivana Mikulic, Lars Dölken, et al. 2013. “CDK8 Kinase Phosphorylates Transcription Factor STAT1 to Selectively Regulate the Interferon Response.” Immunity 38 (2): 250. https://doi.org/10.1016/J.IMMUNI.2012.10.017.

Bodor, Dani L., Mariluz Gómez Rodríguez, Nuno Moreno, and Lars E.T. Jansen. 2012. “Analysis of Protein Turnover by Quantitative SNAP-Based Pulse-Chase Imaging.” Current Protocols in Cell Biology, no. SUPPL.55 (June). https://doi.org/10.1002/0471143030.CB0808S55.

Buenrostro, Jason D., Paul G. Giresi, Lisa C. Zaba, Howard Y. Chang, and William J. Greenleaf. 2013. “Transposition of Native Chromatin for Fast and Sensitive Epigenomic Profiling of Open Chromatin, DNA-Binding Proteins and Nucleosome Position.” Nature Methods 2013 10:12 10 (12): 1213–18. https://doi.org/10.1038/nmeth.2688.

Carpenter, Anne E., Thouis R. Jones, Michael R. Lamprecht, Colin Clarke, In Han Kang, Ola Friman, David A. Guertin, et al. 2006. “CellProfiler: Image Analysis Software for Identifying and Quantifying Cell Phenotypes.” Genome Biology 7 (10): 1–11. https://doi.org/10.1186/GB-2006-7-10-R100/FIGURES/4.

Chavez, Alejandro, Marcelle Tuttle, Benjamin W. Pruitt, Ben Ewen-Campen, Raj Chari, Dmitry Ter-Ovanesyan, Sabina J. Haque, et al. 2016. “Comparison of Cas9 Activators in Multiple Species.” Nature Methods 13 (7): 563–67. https://doi.org/10.1038/NMETH.3871.

Cheon, HyeonJoo, and George R. Stark. 2009. “Unphosphorylated STAT1 Prolongs the Expression of Interferon-Induced Immune Regulatory Genes.” Proceedings of the National Academy of Sciences 106 (23): 9373–78. https://doi.org/10.1073/PNAS.0903487106.

Corces, M. Ryan, Alexandro E. Trevino, Emily G. Hamilton, Peyton G. Greenside, Nicholas A. Sinnott-Armstrong, Sam Vesuna, Ansuman T. Satpathy, et al. 2017. “An Improved ATAC-Seq Protocol Reduces Background and Enables Interrogation of Frozen Tissues.” Nature Methods 2017 14:10 14 (10): 959–62. https://doi.org/10.1038/nmeth.4396.

D’Urso, Agustina, and Jason H. Brickner. 2017. “Epigenetic Transcriptional Memory.” Current Genetics. https://doi.org/10.1007/s00294-016-0661-8.

D’Urso, Agustina, Yoh-hei Takahashi, Bin Xiong, Jessica Marone, Robert Coukos, Carlo Randise-Hinchliff, Ji-Ping Wang, Ali Shilatifard, and Jason H. Brickner. 2016. “Set1/COMPASS and Mediator Are Repurposed to Promote Epigenetic Transcriptional Memory.” ELife 5 (June): e16691. https://doi.org/10.7554/eLife.16691.

Darnell, James E., Ian M. Kerr, and George R. Stark. 1994. “Jak-STAT Pathways and Transcriptional Activation in Response to IFNs and Other Extracellular Signaling Proteins.” Science (New York, N.Y.) 264 (5164): 1415–21. https://doi.org/10.1126/SCIENCE.8197455.

Delgoffe, Greg M., and Dario A.A. Vignali. 2013. “STAT Heterodimers in Immunity: A Mixed Message or a Unique Signal?” JAK-STAT 2 (1): e23060. https://doi.org/10.4161/JKST.23060.

Ding, Yong, Ning Liu, Laetitia Virlouvet, Jean Jack Riethoven, Michael Fromm, and Zoya Avramova. 2013. “Four Distinct Types of Dehydration Stress Memory Genes in Arabidopsis Thaliana.” BMC Plant Biology 13 (1): 1–11. https://doi.org/10.1186/1471-2229-13-229/FIGURES/4.

Dull, Tom, Romain Zufferey, Michael Kelly, R. J. Mandel, Minh Nguyen, Didier Trono, and Luigi Naldini. 1998. “A Third-Generation Lentivirus Vector with a Conditional Packaging System.” Journal of Virology 72 (11): 8463–71. https://doi.org/10.1128/JVI.72.11.8463-8471.1998.

Fanucchi, Stephanie, Ezio T. Fok, Emiliano Dalla, Youtaro Shibayama, Kathleen Börner, Erin Y. Chang, Stoyan Stoychev, et al. 2019. “Immune Genes Are Primed for Robust Transcription by Proximal Long Noncoding RNAs Located in Nuclear Compartments.” Nature Genetics 51 (1): 138–50. https://doi.org/10.1038/s41588-018-0298-2.

Gialitakis, Manolis, Panagiota Arampatzi, Takis Makatounakis, and Joseph Papamatheakis. 2010. “Gamma Interferon-Dependent Transcriptional Memory via Relocalization of a Gene Locus to PML Nuclear Bodies.” Molecular and Cellular Biology 30 (8): 2046–56. https://doi.org/10.1128/MCB.00906-09.

Giménez, C. A., M. Ielpi, A. Mutto, L. Grosembacher, P. Argibay, and F. Pereyra-Bonnet. 2016. “CRISPR-on System for the Activation of the Endogenous Human INS Gene.” Gene Therapy 2016 23:6 23 (6): 543–47. https://doi.org/10.1038/gt.2016.28.

Jeong, Jae Yeon, Hyung Soon Yim, Ji Young Ryu, Hyun Sook Lee, Jung Hyun Lee, Dong Seung Seen, and Sung Gyun Kang. 2012. “One-Step Sequence-and Ligation-Independent Cloning as a Rapid and Versatile Cloning Method for Functional Genomics Studies.” Applied and Environmental Microbiology 78 (15): 5440–43. https://doi.org/10.1128/AEM.00844-12.

Kamada, Rui, Wenjing Yang, Yubo Zhang, Mira C Patel, Yanqin Yang, Ryota Ouda, Anup Dey, et al. 2018. “Interferon Stimulation Creates Chromatin Marks and Establishes Transcriptional Memory.” Proceedings of the National Academy of Sciences of the United States of America 115 (39): E9162–71. https://doi.org/10.1073/pnas.1720930115.

Konermann, Silvana, Mark D. Brigham, Alexandro E. Trevino, Julia Joung, Omar O. Abudayyeh, Clea Barcena, Patrick D. Hsu, et al. 2014. “Genome-Scale Transcriptional Activation by an Engineered CRISPR-Cas9 Complex.” Nature 2014 517:7536 517 (7536): 583–88. https://doi.org/10.1038/nature14136.

Lämke, Jörn, Krzysztof Brzezinka, Simone Altmann, and Isabel Bäurle. 2016. “A Hit-and-Run Heat Shock Factor Governs Sustained Histone Methylation and Transcriptional Stress Memory.” The EMBO Journal 35 (2): 162–75. https://doi.org/10.15252/embj.201592593.

Lau, Colleen M., Nicholas M. Adams, Clair D. Geary, Orr El Weizman, Moritz Rapp, Yuri Pritykin, Christina S. Leslie, and Joseph C. Sun. 2018. “Epigenetic Control of Innate and Adaptive Immune Memory.” Nature Immunology 19 (9): 963. https://doi.org/10.1038/S41590-018-0176-1.

Light, William H., Jonathan Freaney, Varun Sood, Abbey Thompson, Agustina D’Urso, Curt M. Horvath, and Jason H. Brickner. 2013. “A Conserved Role for Human Nup98 in Altering Chromatin Structure and Promoting Epigenetic Transcriptional Memory.” Edited by Tom Misteli. PLoS Biology 11 (3): e1001524. https://doi.org/10.1371/journal.pbio.1001524.

Liu, Ning, and Zoya Avramova. 2016. “Molecular Mechanism of the Priming by Jasmonic Acid of Specific Dehydration Stress Response Genes in Arabidopsis.” Epigenetics & Chromatin 2016 9:1 9 (1): 1–23. https://doi.org/10.1186/S13072-016-0057-5.

Meers, Michael P., Terri D. Bryson, Jorja G. Henikoff, and Steven Henikoff. 2019. “Improved Cut&xrun Chromatin Profiling Tools.” ELife 8 (June). https://doi.org/10.7554/ELIFE.46314.001.

Meers, Michael P., Terri Bryson, and Steven Henikoff. 2019. “A Streamlined Protocol and Analysis Pipeline for CUT&RUN Chromatin Profiling.” BioRxiv, March, 569129. https://doi.org/10.1101/569129.

Migita, Kiyoshi, Atsumasa Komori, Takafumi Torigoshi, Yumi Maeda, Yasumori Izumi, Yuka Jiuchi, Taiichiro Miyashita, Minoru Nakamura, Satoru Motokawa, and Hiromi Ishibashi. 2011. “CP690,550 Inhibits Oncostatin M-Induced JAK/STAT Signaling Pathway in Rheumatoid Synoviocytes.” Arthritis Research and Therapy 13 (3): 1–10. https://doi.org/10.1186/AR3333/FIGURES/8.

Moazed, Danesh. 2011. “Mechanisms for the Inheritance of Chromatin States.” Cell 146 (4): 510–18. https://doi.org/10.1016/J.CELL.2011.07.013.

Mogensen, Trine H. 2018. “IRF and STAT Transcription Factors - From Basic Biology to Roles in Infection, Protective Immunity, and Primary Immunodeficiencies.” Frontiers in Immunology 9 (JAN): 3047. https://doi.org/10.3389/FIMMU.2018.03047.

Nabet, Behnam, Justin M. Roberts, Dennis L. Buckley, Joshiawa Paulk, Shiva Dastjerdi, Annan Yang, Alan L. Leggett, et al. 2018. “The DTAG System for Immediate and Target-Specific Protein Degradation.” Nature Chemical Biology 14 (5): 431. https://doi.org/10.1038/S41589-018-0021-8.

Natoli, Gioacchino, and Renato Ostuni. 2019. “Adaptation and Memory in Immune Responses.” Nature Immunology 2019 20:7 20 (7): 783–92. https://doi.org/10.1038/s41590-019-0399-9.

Netea, Mihai G., Jorge Domínguez-Andrés, Luis B. Barreiro, Triantafyllos Chavakis, Maziar Divangahi, Elaine Fuchs, Leo A.B. Joosten, et al. 2020. “Defining Trained Immunity and Its Role in Health and Disease.” Nature Reviews Immunology. Nature Research. https://doi.org/10.1038/s41577-020-0285-6.

Netea, Mihai G., Jessica Quintin, and Jos W.M. Van Der Meer. 2011. “Trained Immunity: A Memory for Innate Host Defense.” Cell Host & Microbe 9 (5): 355–61. https://doi.org/10.1016/J.CHOM.2011.04.006.

Ni, Zuyao, Elizabeth Karaskov, Tao Yu, Steven M. Callaghan, Sandy Der, David S. Park, Zhaodong Xu, Samantha G. Pattenden, and Rod Bremner. 2005. “Apical Role for BRG1 in Cytokine-Induced Promoter Assembly.” Proceedings of the National Academy of Sciences of the United States of America 102 (41): 14611–16. https://doi.org/10.1073/PNAS.0503070102/SUPPL_FILE/03070FIG12.PDF.

Peignier, Adeline, and Dane Parker. 2020. “Trained Immunity and Host-Pathogen Interactions.” Cellular Microbiology 22 (12): e13261. https://doi.org/10.1111/CMI.13261.

Quintin, Jessica, Sadia Saeed, Joost H.A. Martens, Evangelos J. Giamarellos-Bourboulis, Daniela C. Ifrim, Colin Logie, Liesbeth Jacobs, et al. 2012. “Candida Albicans Infection Affords Protection against Reinfection via Functional Reprogramming of Monocytes.” Cell Host and Microbe 12 (2): 223–32. https://doi.org/10.1016/j.chom.2012.06.006.

Ramana, Chilakamarti V., Moitreyee Chatterjee-Kishore, Hannah Nguyen, and George R. Stark. 2000. “Complex Roles of Stat1 in Regulating Gene Expression.” Oncogene. Nature Publishing Group. https://doi.org/10.1038/sj.onc.1203525.

Ramana, Chilakamarti V., M. Pilar Gil, Robert D. Schreiber, and George R. Stark. 2002. “Stat1-Dependent and -Independent Pathways in IFN-γ-Dependent Signaling.” Trends in Immunology. Elsevier Current Trends. https://doi.org/10.1016/S1471-4906(01)02118-4.

Ramsauer, Katrin, Matthias Farlik, Gordin Zupkovitz, Christian Seiser, Andrea Kröger, Hansjörg Hauser, and Thomas Decker. 2007. “Distinct Modes of Action Applied by Transcription Factors STAT1 and IRF1 to Initiate Transcription of the IFN-Gamma-Inducible Gbp2 Gene.” Proceedings of the National Academy of Sciences of the United States of America 104 (8): 2849–54. https://doi.org/10.1073/PNAS.0610944104.

Rawlings, Jason S., Kristin M. Rosler, and Douglas A. Harrison. 2004. “The JAK/STAT Signaling Pathway.” Journal of Cell Science 117 (8): 1281–83. https://doi.org/10.1242/jcs.00963.

Schroder, Kate, Paul J. Hertzog, Timothy Ravasi, and David A. Hume. 2004. “Interferon-Gamma: An Overview of Signals, Mechanisms and Functions.” Journal of Leukocyte Biology 75 (2): 163–89. https://doi.org/10.1189/JLB.0603252.

Sif, S., A. J. Saurin, A. N. Imbalzano, and R. E. Kingston. 2001. “Purification and Characterization of MSin3A-Containing Brg1 and HBrm Chromatin Remodeling Complexes.” Genes & Development 15 (5): 603–18. https://doi.org/10.1101/GAD.872801.

Siwek, Wojciech, Sahar S.H. Tehrani, João F. Mata, and Lars E.T. Jansen. 2020. “Activation of Clustered IFNγ Target Genes Drives Cohesin-Controlled Transcriptional Memory.” Molecular Cell 80 (3): 396-409.e6. https://doi.org/10.1016/j.molcel.2020.10.005.

Song, Jie, Andrew Angel, Martin Howard, and Caroline Dean. 2012. “Vernalization - a Cold-Induced Epigenetic Switch.” Journal of Cell Science 125 (16): 3723–31. https://doi.org/10.1242/jcs.084764.

Sood, Varun, Ivelisse Cajigas, Agustina D’Urso, William H. Light, and Jason H. Brickner. 2017. “Epigenetic Transcriptional Memory of GAL Genes Depends on Growth in Glucose and the Tup1 Transcription Factor in Saccharomyces Cerevisiae.” Genetics, January, genetics.117.201632. https://doi.org/10.1534/genetics.117.201632.

Sump, Bethany, Donna G Brickner, Agustina D’Urso, Seo Hyun Kim, and Jason H Brickner. 2022. “Mitotically Heritable, RNA Polymerase II-Independent H3K4 Dimethylation Stimulates INO1 Transcriptional Memory.” ELife 11 (May). https://doi.org/10.7554/ELIFE.77646.

Verma, Deepti, Venkata Ramanarao Parasa, Johanna Raffetseder, Mihaela Martis, Ratnesh B. Mehta, Mihai Netea, and Maria Lerm. 2017. “Anti-Mycobacterial Activity Correlates with Altered DNA Methylation Pattern in Immune Cells from BCG-Vaccinated Subjects.” Scientific Reports 2017 7:1 7 (1): 1–10. https://doi.org/10.1038/s41598-017-12110-2.

Wen, Zilong, Zhong Zhong, and James E Darnell. 1995. “Maximal Activation of Transcription by Statl and Stat3 Requires Both Tyrosine and Serine Phosphorylation.” Cell 82 (2): 241–50. https://doi.org/10.1016/0092-8674(95)90311-9.

Zhao, Zuodong, Zhuqiang Zhang, Jingjing Li, Qiang Dong, Jun Xiong, Yingfeng Li, Mengying Lan, Gang Li, and Bing Zhu. 2020. “Sustained Tnf-α Stimulation Leads to Transcriptional Memory That Greatly Enhances Signal Sensitivity and Robustness.” ELife 9 (October): 1–27. https://doi.org/10.7554/eLife.61965.

