## Supplemental Figures for "STAT1 is required to establish but not maintain IFNγ-induced transcriptional memory"

### Supplementary 1

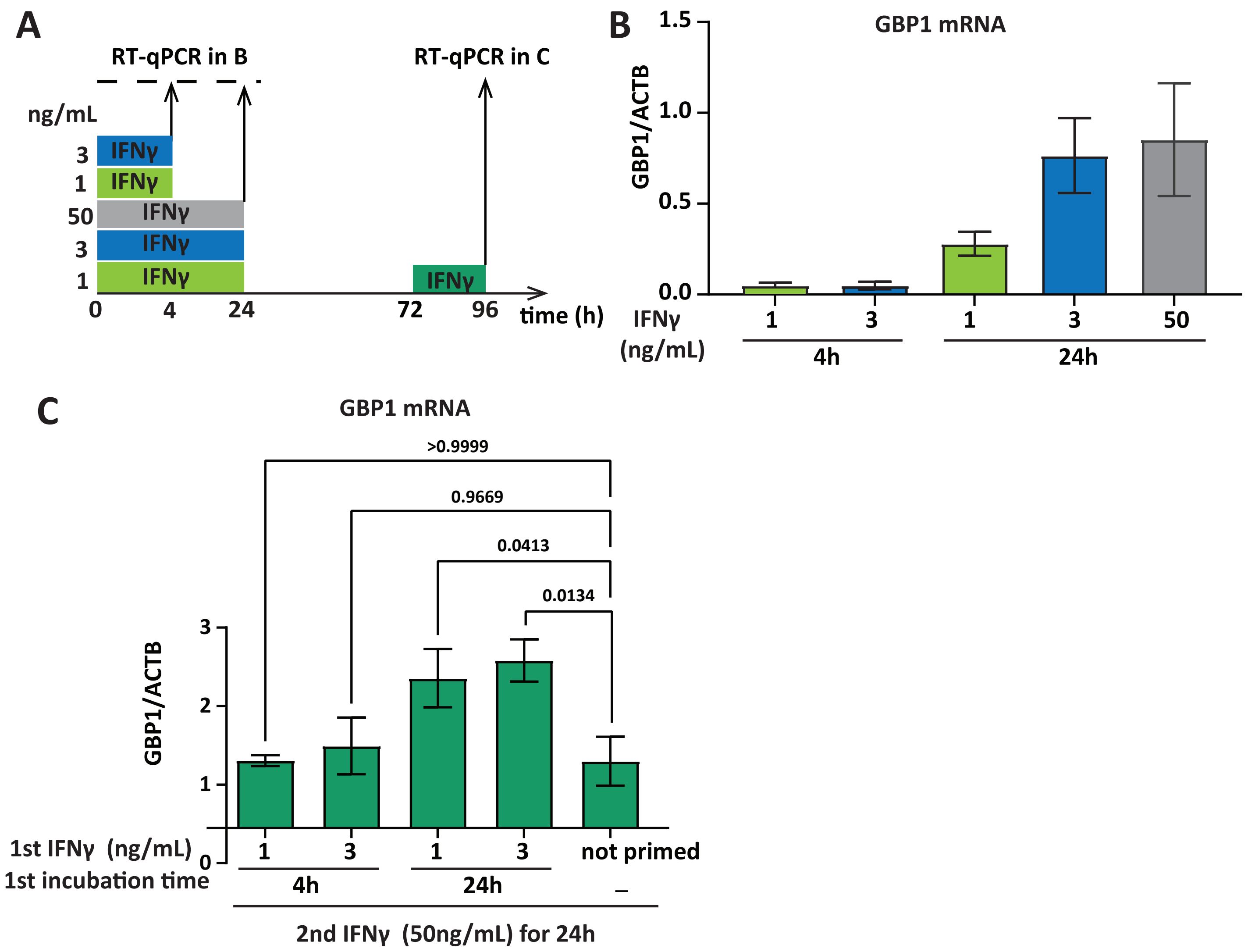

### Supplementary 2

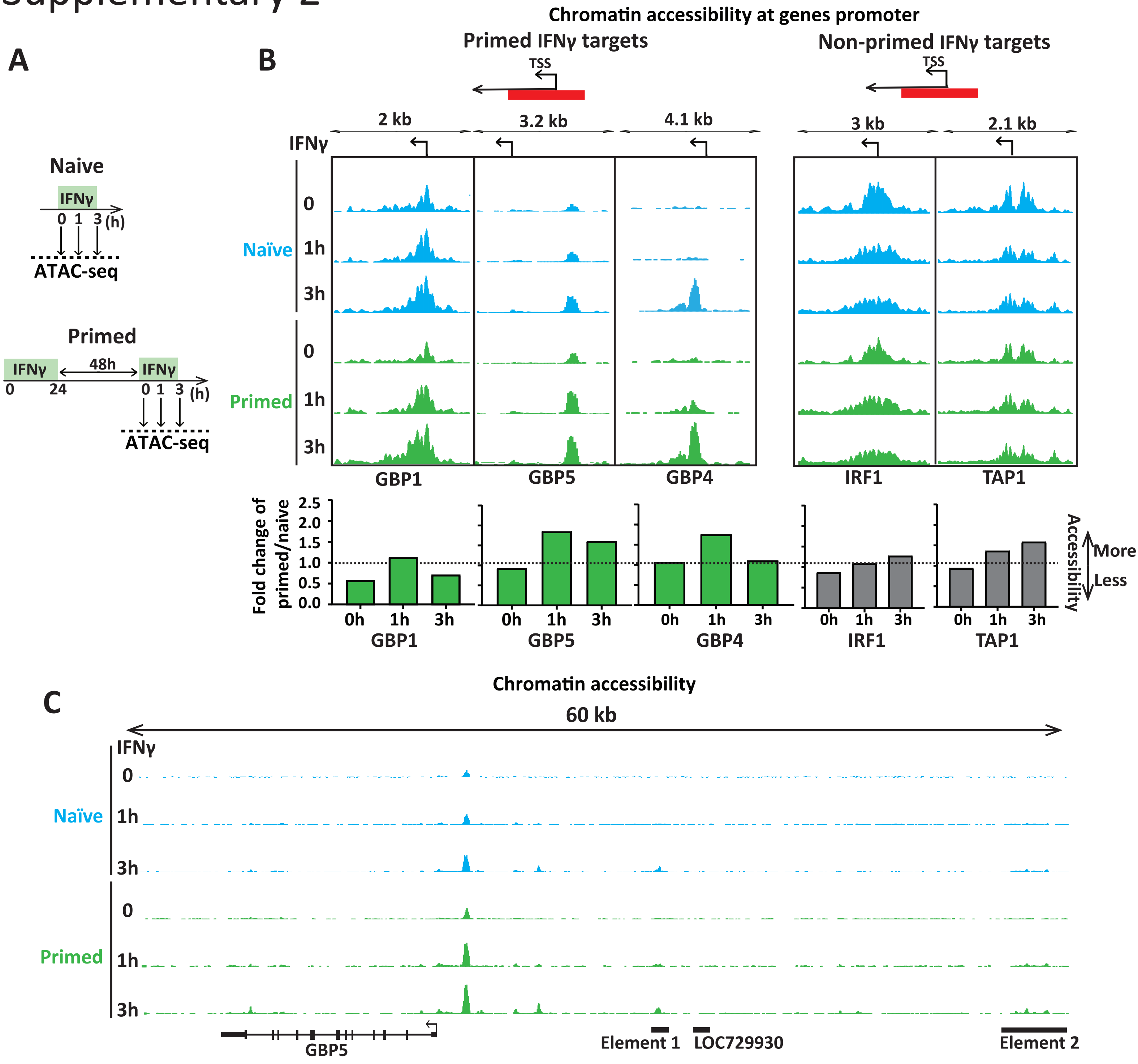

### Supplementary 3

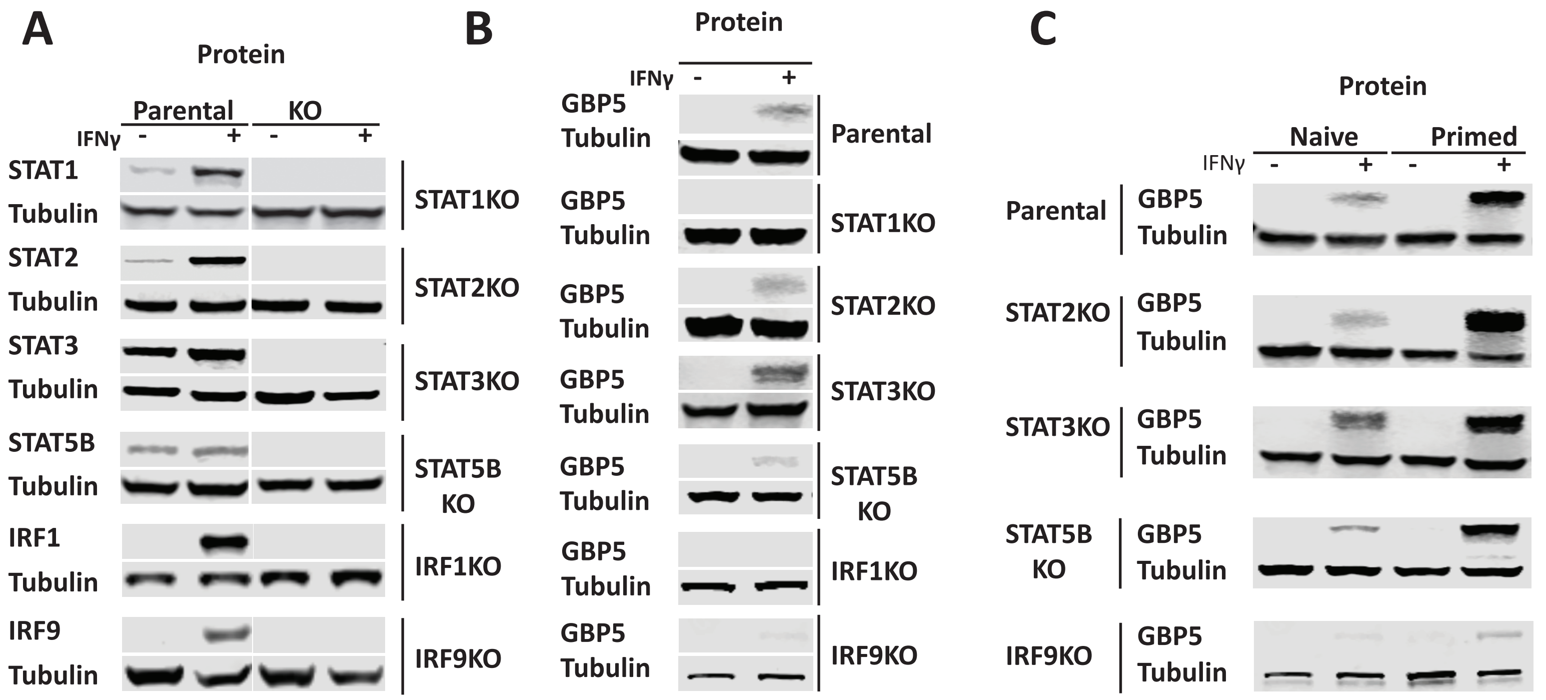

### Supplementary 4

**A**

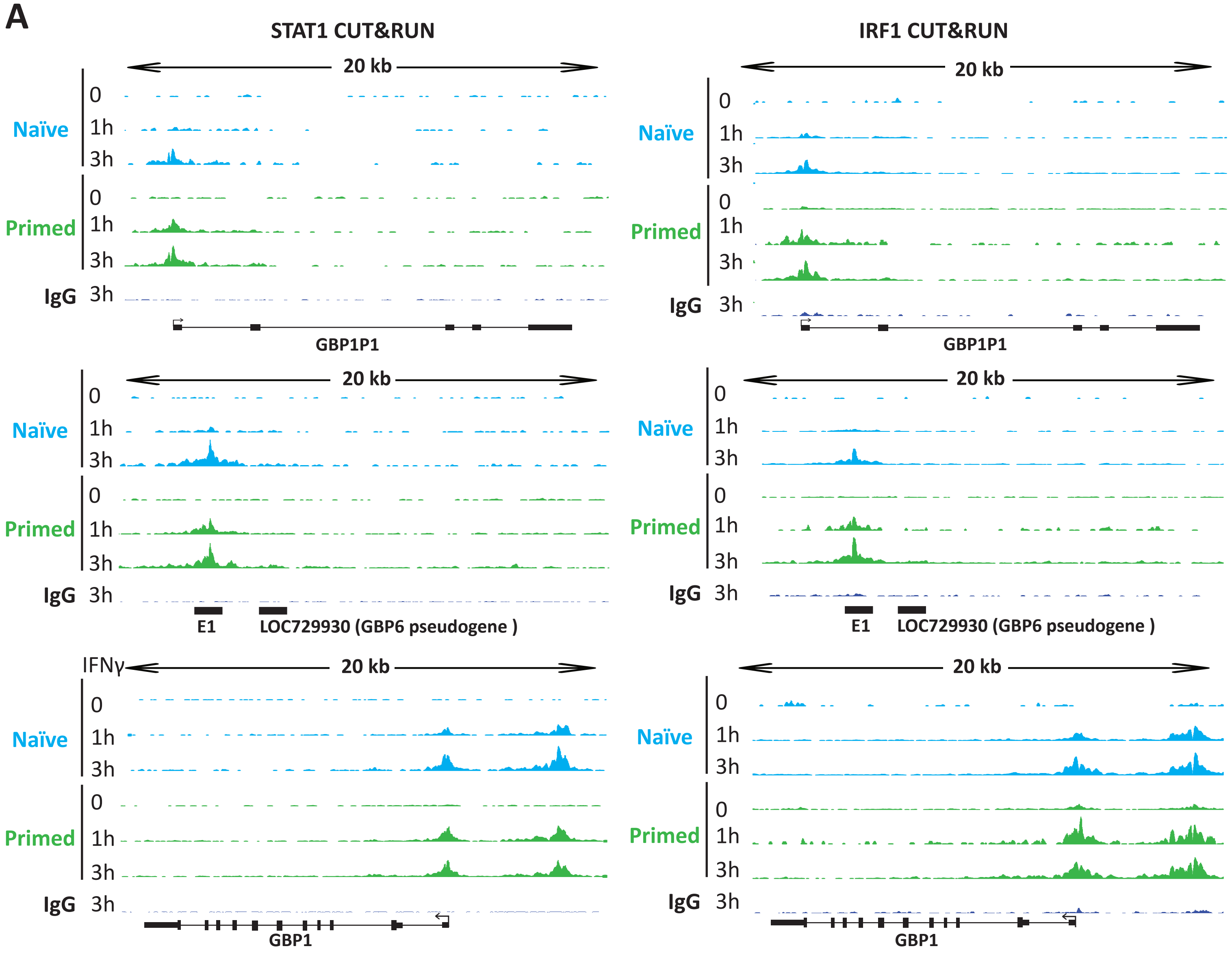


**B** **list of overlap genes in Figure 3G, that showed both faster STAT1 and IRF1 recruitment**

| | Ensembl | Gene name | description |
|---|---|---|---|
| 1 | ENSG00000225492 | GBP1P1 | guanylate binding protein 1 pseudogene |
| 2 | ENSG00000154451 | GBP5 | guanylate binding protein 5 (see Figure 3) |
| 3 | ENSG00000162654 | GBP4 | guanylate binding protein 4 (see Figure 3) |
| 4 | ENSG00000237568 | nan | LncRNA inside GBP5 gene |
| 5 | ENSG00000238081 | LOC729930 | GBP6 pseudogene, transcription factor binding site is found 2.4 kbp upstream of this site |
| 6 | ENSG00000284734 | GBP4 (antisense) | novel transcript, antisense to GBP4 (see Figure 3) |
| 7 | ENSG00000117228 | GBP1 | guanylate binding protein 1 |
| 8 | ENSG00000269588 | LGALS13 | lectin, galactoside-binding, soluble, 13 (LGALS13) pseudogene |
| 9 | ENSG00000253838 | nan | lncRNA inside IDO1 |
| 10 | ENSG00000262151 | CIITA (antisense) | novel transcript, antisense to CIITA |
| 11 | ENSG00000226025 | LGALS17A | galectin 14 pseudogene |
| 12 | ENSG00000131203 | IDO1 | indoleamine 2,3-dioxygenase 1 |

### Supplementary 5

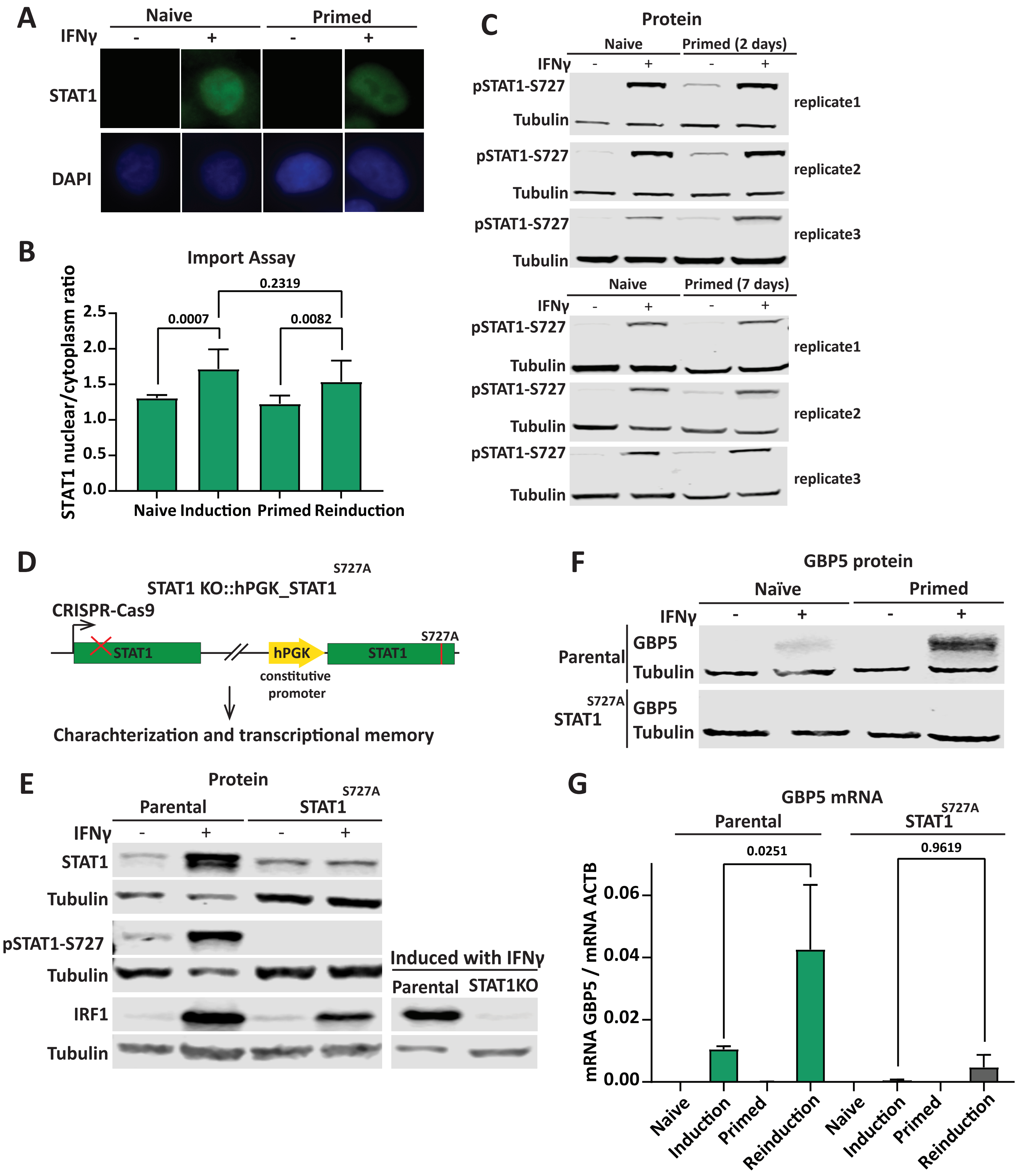

S6. Table1. list of gRNAs and primes.

| **CRISPRa gRNA** | |
|---|---|
| **Primer name #** | **sequence(5'->3')** |
| F_GBP1_SAM_gRNA1 | CACCGAAGCTAAGCAGATTTGTAA |
| F_GBP1_SAM_gRNA2 | CACCGTTCTAAATATTTCATCAATG |
| F_GBP1_SAM_gRNA3 | CACCGAATTAGTGGAGTGTGCCAG |
| F_GBP1_SAM_gRNA4 | CACCGAAATCTTTAAACCCTCCCAC |
| F_GBP1_SAM_gRNA5 | CACCGTTTCACTGTTCCGAAGTTG |
| F_GBP1_SAM_gRNA6 | CACCGTAGAACATGAGTACAACACA |
| R_GBP1_SAM_gRNA1 | AAACTTACAAATCTGCTTAGCTTC |
| R_GBP1_SAM_gRNA2 | AAACCATTGATGAAATATTTAGAAC |
| R_GBP1_SAM_gRNA3 | AAACCTGGCACACTCCACTAATTC |
| R_GBP1_SAM_gRNA4 | AAACGTGGGAGGGTTTAAAGATTTC |
| R_GBP1_SAM_gRNA5 | AAACACAACTTCGGAACAGTGAAAC |
| R_GBP1_SAM_gRNA6 | AAACTGTGTTGTACTCATGTTCTAC |
| F_ASCL1_gRNA1 | CACCGAAGAGGAGGGGGGGGAGTGG |
| F_ASCL1_gRNA2 | CACCGCGGGAGAAAGGAACGGGAGG |
| F_ASCL1_gRNA3 | CACCGCAGCCGCTCGCTGCAGCAG |
| R_ASCL1_gRNA1 | AAACCCACTCCCCCCCCTCCTCTTC |
| R_ASCL1_gRNA2 | AAACCCTCCCGTTCCTTTCTCCCGC |
| R_ASCL1_gRNA3 | AAACCTGCTGCAGCGAGCGGCTGC |
| **CRISPR/Cas9** | |
| **Primer name #** | **sequence(5'->3')** |
| IRF1B_F | CACCGCATGGCTGGGACATCAACA |
| IRF9B_F | CACCGAGGGCTCAGCAACATCCATG |
| STAT1B_F | CACCGAGAACACGAGACCAATGGTG |
| STAT2B-F | CACCGAAGAATAGCATGGTAGCCT |
| STAT3A-F | CACCGCTACAGTGACAGCTTCCCAA |
| STAT5BA-F | CACCGTGCGGCATTATTTATCCCAG |
| ContorlA-F | CACCGTATTACTGATATTGGTGGG |
| ControlB-F | CACCGTTCCGCGTTACATAACTTA |
| IRF1B_R | AAACTGTTGATGTCCCAGCCATGC |
| IRF9B_R | AAACCATGGATGTTGCTGAGCCCTC |
| STAT1B_R | AAACCACCATTGGTCTCGTGTTCTC |
| STAT2B-R | AAACAGGCTACCATGCTATTCTTC |
| STAT3A-R | AAACTTGGGAAGCTGTCACTGTAGC |
| STAT5BA-R | AAACCTGGGATAAATAATGCCGCAC |
| ContorlA-R | AAACCCCACCAATATCAGTAATAC |
| ControlB-R | AAACTAAGTTATGTAACGCGGAAC |
| **qPCR** | |
| Primer name # | sequence(5'->3') |
| GBP1_qPCR_F | GTGGAACGTGTGAAAGCTGA |

| | |
|---|---|
| GBP1_qPCR_R | CAACTGGACCCTGTCGTTCT |
| GBP5_trancC_F | TTCAATTTGCCCCGTCTGTG |
| GBP5_trancC_R | AGGCAGTGTTTCAAGTTGGG |
| b-actin Forward | AACTGGAACGGTGAAGGTGACAGC |
| b-actin Reverse | TGGCTTTTAGGATGGCAAGGGAC |
